# Two Amazon freshwater sponges, *Drulia brownii* and *Tubella paulula*, share a resilient microbiome strongly shaped by seasonality and urbanization

**DOI:** 10.64898/2026.07.30.741787

**Authors:** Lucas S. Silva, James Leão de Araújo, Graciene do Socorro Taveira Fernandes, Rafael Azevedo Baraúna, Diego Assis das Graças, Artur Silva, Maria Paula Cruz Schneider

## Abstract

Sponges are among the earliest-diverging metazoans, and their evolutionary success has been strongly linked to their symbiosis with microorganisms. While marine sponge-microbiome associations have been extensively characterized, freshwater sponges remain comparatively understudied, particularly in tropical systems such as the Amazon, where sponges undergo seasonal flood pulses and urbanization-derived disturbance. Here we characterized the microbiomes of two freshwater sponge species, *Drulia brownii* and *Tubella paulula*, from two contrasting sites in the Tapajós River (Amazon basin), a non-urbanized and an urbanized site, at two time points, the rainy and dry seasons. The whole metagenome was sequenced using a long-read shotgun approach, and metabarcoding was employed to characterize the gemmule microbiome. The analyses revealed that microbiome composition was primarily influenced by season, with urbanization exerting a secondary but significant effect. In non-urbanized rainy-season sponges, the microbiome was dominated by Pseudomonadales and Bacillales, whereas urbanized sponges showed more diverse profiles enriched with Burkholderiales, a possible symbiont. During the dry season, communities converged across sites, with Burkholderiales becoming the dominant taxon, including in gemmules. The 23 high-quality MAGs recovered revealed symbiosis-related genes, broad biosynthetic repertoires, and a distinctive carbohydrate-active enzyme profile. Functional analyses suggest that Pseudomonadales and Burkholderiales may play complementary roles across seasons, and Bacillales may be associated with organic matter turnover. These results show that Amazonian freshwater sponges harbor stress-sensitive but resilient microbiomes, with seasonality driving major compositional shifts and urbanization accelerating convergence toward Burkholderiales-dominated consortia, and highlight the central role of bacterial symbionts in nutrient acquisition, photoprotection, and chemical defense.

## INTRODUCTION

Sponges (phylum Porifera) are filter-feeding, sessile, and ancient macroinvertebrates that emerged approximately 600 million years ago ^1^. Among the earliest diverging metazoans ^2,3^, these are ubiquitous animals that can be found in almost all aquatic habitats: marine sponges account for more than 9,400 species ^4^, comprising environments like deep sea ^5^, bays ^6^ and coral reefs ^7^; however, one of the main evidence of their evolutionary success and adaptability was their colonization and widespread across freshwater environments ^8^, accounting more than 270 valid species ^4^ commonly found in lentic and lotic waters, like rivers, streams, ponds, lakes and caves, and associated to natural or human made structures ^9^. Freshwater sponges (order Spongillida; family Demospongiae) are present across all continents except Antarctica and exhibit high endemism, with a few known cosmopolitan species, such as *Spongilla lacustris*, *Ephydatia fluviatilis*, and *Ephydatia muelleri* ^10,11^.

The evolutionary success of sponges is especially associated to their symbiotic association with microorganisms ^12^, since bacteria may have provided nutritional support since their early existence ^13^. Given their consortium-based nature, sponges are considered complex holobionts ^14,15^, and microorganisms make up to 38% of their biomass ^16^, supporting the health of sponges through supplying mainly carbon sources and nitrogen, and chemical defense ^17–20^. While marine sponges are well-described organisms that play a crucial role in benthic environments, serving as protagonists in the transfer of organic matter to higher trophic levels ^21–24^ and nutrient recycling, their microbiome is a key element in this process ^15^.

Most of the studies on the microbiome of sponges focused on marine species, which have been extensively studied ^25,26^ and are described as highly stable ^27,28^ and actively selected by the host ^29,30^. The microbiome of freshwater sponges, however, remains poorly understood. A few microbiome studies have been conducted for freshwater sponges, most of them with sponges from the highly stable ancient Lake Baikal ^31–34^ or focusing on the cosmopolitan species ^11,35^, with evidence for an actively selected ^36^ and vertically transmitted ^37^ microbiome. For tropical freshwater sponges, a first microbiome study conducted in *Tubella variabilis* (Spongillidae) showed them harboring a microbiome richer than marine sponges ^38^, while another work describing the gemmule-associated microbiome of a *Metania reticulata* (Metaniidae) found a microbiome dominated by unclassified bacteria and Gammaproteobacteria ^39^.

Amazon is a hotspot of diversity and harbors a large diversity of freshwater sponges. However, there is a gap in understanding the microbiome of Amazon sponges, as well as how seasonal changes in the Amazon and human activity affect it. Here, a metagenomic approach was used to characterize the microbiome associated with two Amazon freshwater sponges, *Drulia brownii* and *Tubella paulula*, to understand how Amazon seasonality and urbanization influence the microbial community of the sponges and their gemmules, and to assess the effects of long-term exposure to the dry season. A metagenome-resolved genomics approach was used to identify key metabolic pathways in sponge-associated bacterial consortia across seasons, providing insights into how the sponge holobiont adapts to obtain carbon sources, nutrients, and protection.

## METHODS

### Experimental design, sampling strategy, and morphological identification

Sampling was conducted in the Tapajós River (Santarém, Pará, Brazil), a clear-water system characterized by low turbidity, reduced dissolved and particulate organic matter, and oligotrophic conditions ^40^, which is highly influenced by Amazonian seasonality (Figure 1a-e). Two contrasting sites were selected: (i) Praia do Cajueiro (2°30’04.8”S 54°57’26.2”W), an anthropogenically impacted freshwater beach subject to intense urbanization and used as bathing waters by the population, (ii) Ponta do Cururu (2°27’55.1”S 54°58’40.5”W), a non-urbanized site about 4.7 km away from Praia do Cajueiro, accessed by boat (Figure 1c). Sampling campaigns were conducted during two distinct hydrological extremes of the Amazon flood pulse: the rainy season, characterized by high water levels in July 2023 (Figure 1e), and the dry season, marked by low water levels in November 2023 (Figure 1d).

**Figure 1.**
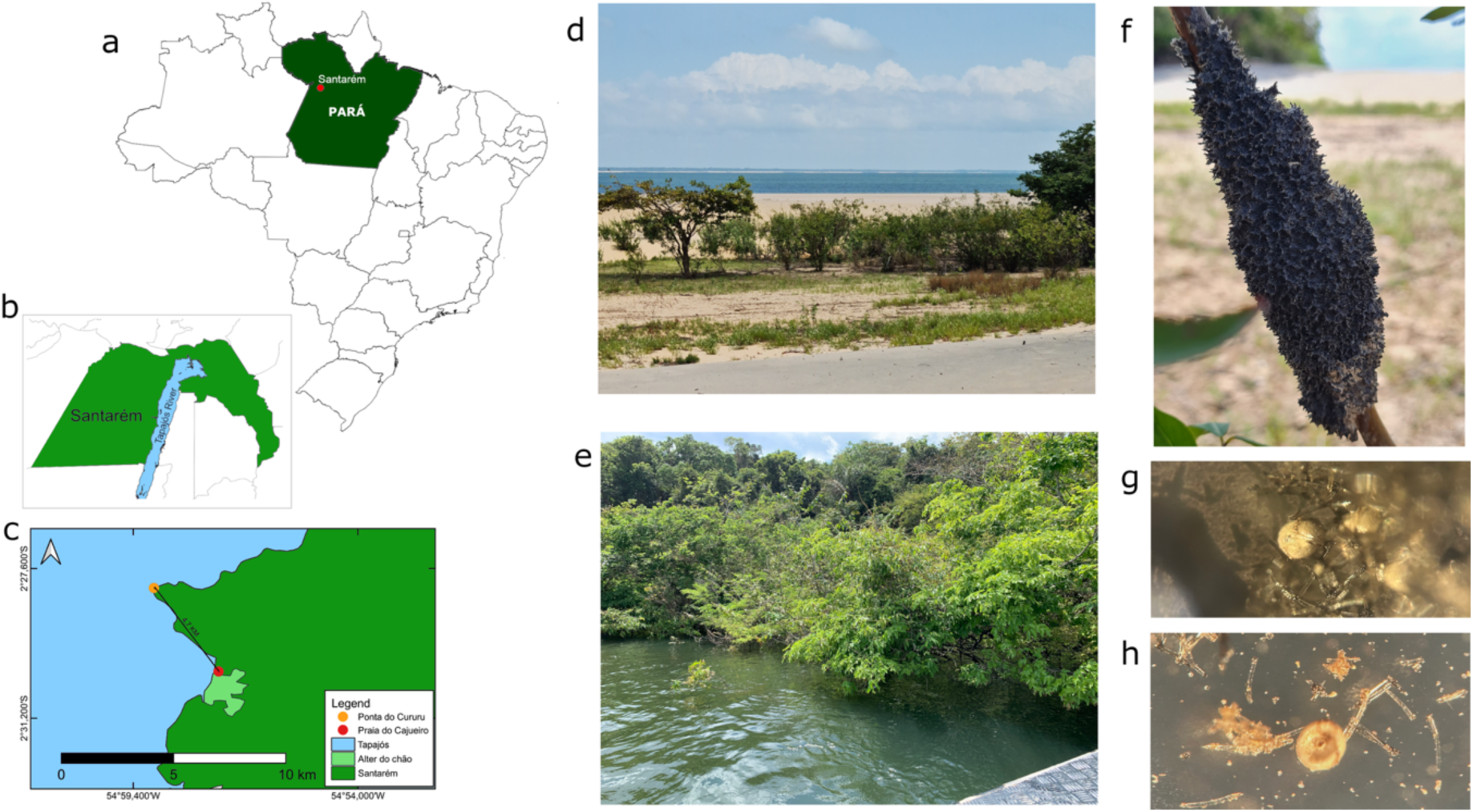
Study area and freshwater sponge samples from the Tapajós River (Pará, Brazil). The figure shows (a) a map of Brazil highlighting the state of Pará in green, with the municipality of Santarém indicated by a red dot; (b) Map of Santarém municipality (green area) highlighting the Tapajós River in blue; (c) Sampling sites along the Tapajós River at the district of Alter do Chão (green area), within Santarém: i) Ponta do Cururu (orange dot, non-urbanized site) and ii) Praia do Cajueiro (red dot, urbanized site); (d) Ponta do Cururu during the dry season (November 2023); (e) Ponta do Cururu during the rainy season (July 2023); (f) *D. brownii* collected at Ponta do Cururu during the dry season; (g-h) Gemmules of Amazon freshwater sponges observed under a stereomicroscope.

A total of 39 sponge specimens were collected. Morphological identification was carried out using standard spicule and gemmule preparation protocol: approximately 0.5 cm fragments of sponge tissue were digested in 35% hydrogen peroxide at 65 °C until complete organic dissolution, then washed, centrifuged, and used for slide preparation for light microscopy analysis, following a method and taxonomy guide already described ^41^, and confirmed using genus-specific works ^42,43^. For specimens selected for the metagenome, approximately 50 gemmules per specimen were hand-picked under a stereomicroscope (Nikon C-DSD115) to ensure accurate genus-level identification by examining gemmoscleres (Figure S1). Subsequently, 100 gemmules per specimen were manually isolated for DNA isolation and metabarcoding sequencing (Figure 1e-f). Microscopic analyses were performed on a Leica DM750 microscope equipped with a Flexacam i5 digital imaging system.

### DNA isolation, COI barcoding, and phylogenetic analysis

DNA was isolated from bulk sponge tissue and also from sponge gemmules using the FastDNA SPIN Kit for Soil (MP Biomedicals), following the manufacturer’s protocols. Given the importance of the epibacterial layer for freshwater sponge gemmules’ healthy hatching ^37^, gemmules were not sterilized before DNA isolation. DNA quality was assessed on 1% agarose gels.

Taxonomic validation through molecular identification was performed by amplifying the cytochrome c oxidase subunit I (COI) gene using 100 µL PCR reactions with GoTaq Green Master Mix (Promega) for the 39 samples using the primers LCO1490 (GGTCAACAAATCATAAAGATATTGG) and HCO2198 (TAAACTTCAGGGTGACCAAAAAATCA) ^44^, each at a final concentration of 0.1 µM/uL and 0.2 ng/uL of sponge DNA as template. Amplicons were cloned into pGEM-T Easy vectors (Promega) and transformed into *E. coli* DH10B, and eight clones per specimen were sequenced by Sanger method on an ABI 3500 Genetic Analyzer (Thermo Fisher) with BigDye Terminator v3.1 (Thermo Fisher). Raw ABI sequences were manually curated to trim low-quality extremities using the software Sequencher v4.1.4.

Sequences were aligned with reference freshwater sponge sequences from GenBank (see Table S1) using MAFFT v7 online server (https://mafft.cbrc.jp/alignment/server/) ^45^. Alignments were checked in BioEdit v7.0.5 ^46^, and the phylogenetic reconstruction was conducted in MEGA7 ^47^ using the GTR+G substitution model (the best selected by MEGA7), with 1000 bootstrap replicates to assess node support. The phylogeny was visualized with ggtree (R package) and used to define representative samples for metagenomic analyses while minimizing redundant clonal specimens.

To characterize microbial communities within gemmules, PCR amplification of bacterial 16S rRNA was performed using the universal primers forward 27F (AGAGTTTGATCMTGGCTCAG) and reverse 1492R (TACGGYTACCTTGTTACGACTT) primers, as performed for eukaryotic 18S rRNA genes (8F GAAACTGCGAATGGCTC and 1520R CYGCAGGTTCACCTAC). PCR reactions were performed using GoTaq Colorless Master Mix (Promega) in 100 µL reactions, with 0.2 ng/uL of the gemmule-obtained DNA as template and primers at a final concentration of 0.1 µM/uL. Some PCRs didn’t yield the minimum amplicon amount necessary for sequencing, despite the experimental efforts.

### Shotgun metagenome and amplicon sequencing

For the whole metagenome, two sponge species were selected that were consistently represented across samples. One species was sampled in both urbanized and non-urbanized sites, whereas the second species was restricted to the non-urbanized site, as it was not detected in the impacted area. Both species were collected in two contrasting hydrological seasons (rainy and dry seasons), and for each species-condition combination, three biological replicates were included. This design yielded six experimental conditions (2 species × 2 seasons × site representation) and 18 metagenomic libraries, ensuring replication across seasonal and disturbance gradients.

Selected specimens were subjected to shotgun sequencing on a PromethION P2 platform (Oxford Nanopore Technologies). Sequencing libraries were prepared using the Ligation Sequencing kit for gDNA, with barcoding performed using the Native Barcoding SQK-NBD114.96 for R10.4.1 flow cells. Shotgun sequencing was performed in four independent runs to reach a minimum target yield of 3 Gb per specimen. The gemmules 16S and 18S amplicons were also sequenced on the PromethION P2 using the Ligation Sequencing Amplicon kit and following ONT amplicon sequencing workflows, with a minimum yield threshold of 70,000 reads per library. Raw data were basecalled with Dorado (ONT), and reads with a length <1 kb and Phred score <12 were discarded using NanoFilt v2.8.0 ^48^.

### Metagenomic profiling, assembly, and genome binning

Two analytical pipelines were employed for the high-quality reads. Taxonomic profiling of both shotgun and amplicon datasets was conducted with Metaxa2 v2.2.3 ^49^ under default configurations for bacteria and eukaryotes, based on the 16S and 18S databases, respectively. Two assembly strategies were employed. For genome-based metagenomic analysis, triplicate samples from the same condition were co-assembled into a single dataset, enabling broader recovery of low-abundance genomic fragments. For genome-resolved metagenomics, assemblies were carried out individually for each sample, ensuring that metagenome-assembled genomes (MAGs) could be traced back to specific specimens. Both strategies used Flye v2.9.5 ^50^ to retrieve high-quality contigs. For the latter, binning was performed through an initial mapping of high-quality reads against the individual metagenome contigs using minimap2 v2.28 ^51^, followed by the conversion of the SAM alignment file to a sorted BAM format, which was used as input, along with metagenomic contigs, for semibin2 v2.2.0 ^52^. The retrieved MAGs were assigned taxonomically and evaluated for completeness and contamination using CheckM v1.2.3 ^53^. Those with completeness >70% and contamination <10% were used for dereplication, which was conducted with dRep v3.6.2 ^54^. All the dereplicated MAGs retained for further analysis showed >76.7% completeness and <6.7% contamination. From an initial set of 43 bins (see Table S2), 23 high-quality MAGs were selected (Figure A3, Table 1). For MAGs species-level identification, the 16S rRNA genes were retrieved from high-quality MAGs using Barrnap v0.9 (https://github.com/tseemann/barrnap) and underwent BLAST against the Genbank nucleotide database.

**Table 1.**
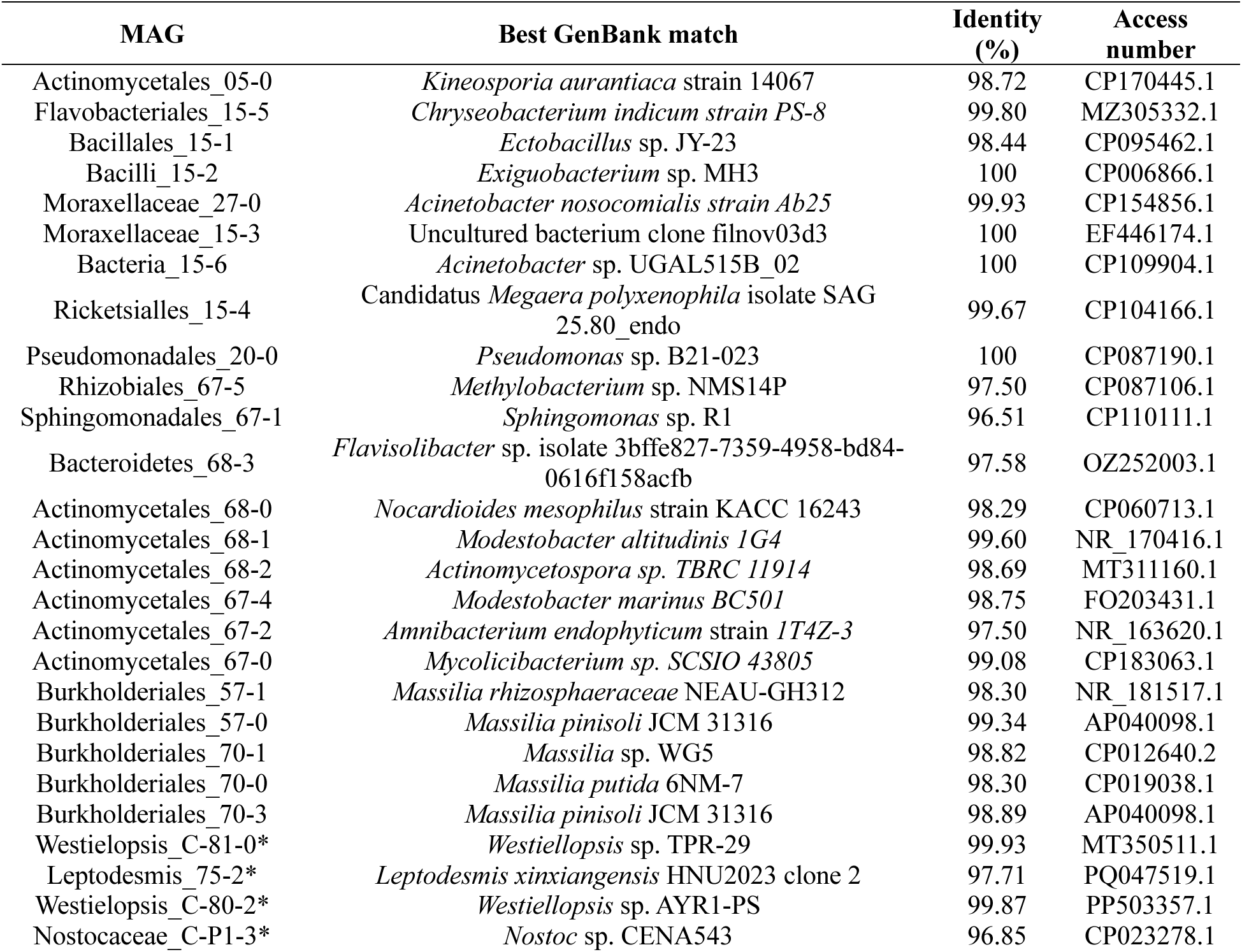
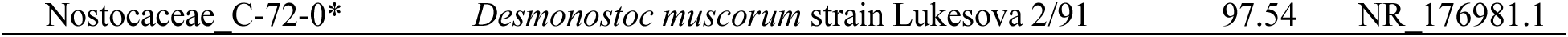
Dereplicated Metagenome-Assembled Genomes retrieved from Amazon freshwater sponges. All the genomes shown are labeled according to the CheckM classification (MAG column) and show the closest strain identified by 16S-based BLAST using NCBI GenBank as the reference. Genomes marked with an asterisk (*) were retrieved through cultivation methods.

| MAG | Best GenBank match | Identity (%) | Access number |
| --- | --- | --- | --- |
| Actinomycetales_05-0 | *Kineosporia aurantiaca* strain 14067 | 98.72 | CP170445.1 |
| Flavobacteriales_15-5 | *Chryseobacterium indicum* strain PS-8 | 99.80 | MZ305332.1 |
| Bacillales_15-1 | *Ectobacillus* sp. JY-23 | 98.44 | CP095462.1 |
| Bacilli_15-2 | *Exiguobacterium* sp. MH3 | 100 | CP006866.1 |
| Moraxellaceae_27-0 | *Acinetobacter nosocomialis* strain Ab25 | 99.93 | CP154856.1 |
| Moraxellaceae_15-3 | Uncultured bacterium clone filnov03d3 | 100 | EF446174.1 |
| Bacteria_15-6 | *Acinetobacter* sp. UGAL515B_02 | 100 | CP109904.1 |
| Rickettsiales_15-4 | Candidatus *Megaera polyxenophila* isolate SAG 25.80_endo | 99.67 | CP104166.1 |
| Pseudomonadales_20-0 | *Pseudomonas* sp. B21-023 | 100 | CP087190.1 |
| Rhizobiales_67-5 | *Methylobacterium* sp. NMS14P | 97.50 | CP087106.1 |
| Sphingomonadales_67-1 | *Sphingomonas* sp. R1 | 96.51 | CP110111.1 |
| Bacteroidetes_68-3 | *Flavisolibacter* sp. isolate 3bffe827-7359-4958-bd84-0616f158acfb | 97.58 | OZ252003.1 |
| Actinomycetales_68-0 | *Nocardioide mesophilus* strain KACC 16243 | 98.29 | CP060713.1 |
| Actinomycetales_68-1 | *Modestobacter altitudinis* IG4 | 99.60 | NR_170416.1 |
| Actinomycetales_68-2 | *Actinomycetospira* sp. TBRC 11914 | 98.69 | MT311160.1 |
| Actinomycetales_67-4 | *Modestobacter marinus* BC501 | 98.75 | FO203431.1 |
| Actinomycetales_67-2 | *Amnibacterium endophyticum* strain IT4Z-3 | 97.50 | NR_163620.1 |
| Actinomycetales_67-0 | *Mycolicibacterium* sp. SCSIO 43805 | 99.08 | CP183063.1 |
| Burkholderiales_57-1 | *Massilia rhizosphaeraceae* NEAU-GH312 | 98.30 | NR_181517.1 |
| Burkholderiales_57-0 | *Massilia pinisoli* JCM 31316 | 99.34 | AP040098.1 |
| Burkholderiales_70-1 | *Massilia* sp. WG5 | 98.82 | CP012640.2 |
| Burkholderiales_70-0 | *Massilia putida* 6NM-7 | 98.30 | CP019038.1 |
| Burkholderiales_70-3 | *Massilia pinisoli* JCM 31316 | 98.89 | AP040098.1 |
| Westiellopsis_C-81-0* | *Westiellopsis* sp. TPR-29 | 99.93 | MT350511.1 |
| Leptodesmis_75-2* | *Leptodesmis xinxiangensis* HNU2023 clone 2 | 97.71 | PQ047519.1 |
| Westiellopsis_C-80-2* | *Westiellopsis* sp. AYR1-PS | 99.87 | PP503357.1 |
| Nostocaceae_C-P1-3* | *Nostoc* sp. CENA543 | 96.85 | CP023278.1 |

### Functional annotation and pathways reconstruction analyses

Functional annotation for both whole metagenome assemblies and high-quality MAGs was carried out using eggNOG-mapper v2.1.12 with the eggNOG database 5.0.2 ^55^, which provided the assignment of orthologous groups and the reconstruction of KEGG pathways, BRITE hierarchies, and KEGG modules through the KEGG Mapper - Resconstruct web tool (https://www.genome.jp/kegg/mapper/reconstruct.html). This allowed the inference of broad metabolic potential across samples. To refine the functional interpretation of secondary metabolism, biosynthetic gene clusters (BGCs) were predicted using antiSMASH v7.0 ^56^ under relaxed parameters. All predicted clusters were subsequently manually inspected and curated to ensure accuracy.

Carbohydrate-active enzymes (CAZymes) were identified by MAGs against the CAZy database using the dbCAN v2.0.11 with the database dbCAN-HMMdb-V13 (https://bcb.unl.edu/dbCAN2/), enabling the characterization of enzymatic repertoires related to carbohydrate degradation and modification. In addition, genomic determinants of antiphage and antiplasmid defense systems were detected with DefenseFinder v2.0.0 ^57^ under default configurations, allowing the quantification of defense-associated loci per MAG.

### Statistical analysis

For taxonomic profiles, only the most abundant taxa were retained for visualization: the 15 most abundant orders for bacteria and the 15 most abundant classes for eukaryotes. Relative abundances were calculated after normalization, and group-level summaries were used to construct stacked barplots. Correlation analyses between microbial taxa and metadata variables (season and urbanization) were assessed using Pearson’s correlation coefficient, and significance was determined after false discovery rate (FDR) correction (Benjamini-Hochberg), with thresholds set at p < 0.05, p < 0.01, and p < 0.001. Alpha diversity metrics (Richness, Shannon index, and Pielou’s evenness) were computed per sample using the vegan R package, and differences across conditions were visualized as boxplots.

Community-level beta diversity was assessed using distance-based ordination methods tailored to the type of data analyzed. For taxonomic composition, including bacterial and eukaryotic community profiles derived from 16S and 18S rRNA genes, dissimilarities were calculated with the Bray-Curtis index, and ordinations were performed by non-metric multidimensional scaling (NMDS). NMDS was also employed for compositional patterns of biosynthetic gene clusters (BGCs) and defense system repertoires. KEGG pathway presence/absence and CAZyme family profiles were analyzed by principal coordinates analysis (PCoA), where genome-level gene abundances provided the primary signal. Group-level separation in ordination space was statistically tested by PERMANOVA (999 permutations) and included 95% confidence ellipses.

Functional profiles (KEGG pathways, biosynthetic gene clusters, carbohydrate-active enzymes, and defense systems) were summarized as count matrices per genome or per pathway. Differences in abundance across groups were tested using Wilcoxon rank-sum tests, with p-values adjusted by Benjamini-Hochberg correction, and results were considered significant at adjusted p < 0.05. Differential abundance analyses of eukaryotic and bacterial taxa were reported as log₂ fold changes against the background distribution, with adjusted significance thresholds applied. All statistical analyses and visualizations were conducted in R v4.3.1.

## RESULTS

We first performed morphological characterization of all sponge samples obtained, given the limited molecular information available for non-cosmopolitan freshwater sponges, and cloned the COI gene to sequence and select only non-ambiguous samples since sponges are known for epibiotic structured assemblies. Then, 18 samples were selected for long-read shotgun metagenomic sequencing, comprising two contrasting sites from both the rainy and dry Amazon seasons; for *Drullia brownii*, metabarcoding was employed to sequence the gemmule microbiome. Additionally, we also attempted to cultivate possible symbiont lineages. Our morphological analysis revealed that the 39 samples used in this study belonged primarily to two neotropical sponge genera: *T. paulula* (family Spongillidae) and *D. brownii* (family Metaniidae). To understand their phylogenetic placement among other freshwater sponges, a COI-based phylogenetic analysis was conducted. The results successfully distinguished the two families identified through morphology and revealed that the sampled Amazon sponges belonged to three distinct clades (Figure S2). The first clade (Figure S2, clade A) grouped all *D. brownii* sponges, the most represented genus among samples, into a monophyletic group. Interestingly, the closest clade (Figure S2, clade B) to *D. brownii* was a *Corvospongilla ultima* (Spongillidae) sample from India (Bolotov, 2018), which grouped with a sample labeled as *D. brownii* but containing both *Drulia* and *Corvospongilla* gemmoscleres, a probable epibiotic event, known to occur in sponges (Nunes, 2024). Both clades (Figure S2, clades A and B) were found near the *Corvomeyenia* sp. Addis 03-32 (Metaniidae) clade and the tree root. The third clade (Figure S2, clade C) contains some of the specimens clustered with *T. variabilis* samples from Brazil’s northwest region (Laport et al., 2019), within the family Spongillidae. These molecular results confirmed the morphological findings and guided us in selecting samples for whole microbiome shotgun sequencing.

### *D. brownii* and *T. paulula* share a dynamic microbiome shaped by stressful conditions

Whole metagenomic sequencing of the 18 bulk sponge samples enabled recovery of microbial community structure using a metabarcoding approach encompassing both eukaryotes and prokaryotes. The overall community (prokaryotic and eukaryotic) composition of the Amazon freshwater sponge’s consortium showed a highly similar profile for both sponge species, *D. brownii* and *T. paulula*, with no evidence for a taxonomy-specific microbial community. Overall, it was mainly composed of bacteria, which accounted for more than 50% of the community across the samples and conditions studied in this work, reaching 75% in non-urbanized sites (Figure S3).

The prokaryotic composition of the sponge microbiome was heterogeneous among individuals, as expected for sponges given the strength of host-microbiome interactions (Thomas et al., 2016), and was primarily dominated by Betaproteobacteria, Gammaproteobacteria, and Bacilli, and exhibited considerable differences across the sampled conditions (Figure 2a). During the rainy season, sponges exhibited distinct microbial profiles, with notable differences in microbial composition between sites highly affected by urbanization (Praia do Cajueiro) and the less affected (Ponta do Cururu) (Figure 2c), since the urbanized site produced a more even profile on the *D. brownii* microbiome (Figure S3). Regarding bacterial composition, sponges from non-urbanized sites were dominated by two bacterial orders, Pseudomonadales and Bacillales (Figure 2a). Conversely, samples from urbanized sites showed a significant shift, with diversification of bacterial composition, reductions in Pseudomonadales and Bacillales, and an increase in Burkholderiales, which were minorly represented in non-urbanized conditions.

**Figure 2.**
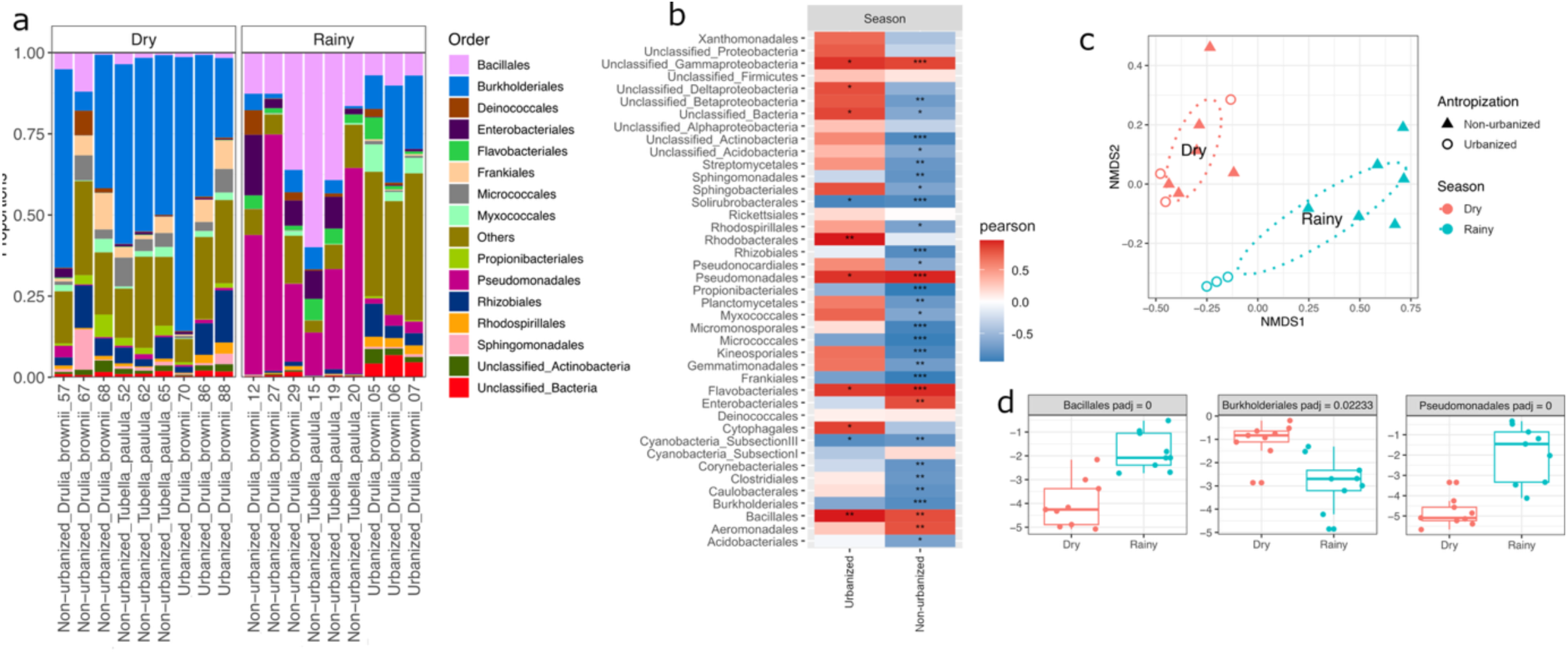
Bacterial diversity of Amazonian sponges across rainy and dry seasons, and for urbanized and non-urbanized sites. Figure (a) shows the relative abundance of the 15 main prokaryotic taxa at the order level across sponge samples grouped per conditions; (b) correlations between the 41 most abundant taxa and the variable season for urbanized and non-urbanized sites. The colors indicate Pearson correlation direction and strength, with blue indicating negative, red for positive, and asterisks showing statistically significant correlations after FDR correction (* p < 0.05, ** p < 0.01, *** p < 0.001); (c) NMDS of prokaryotic composition profile based on Bray-Curtis dissimilarity. Points represent individual samples, colored by season and shaped by site condition. Dotted ellipses denote 95% confidence intervals for seasonal groups; and (d) boxplots showing the log-relative normalized abundance of selected taxa found to be significantly different between seasons (padj < 0.05). Colors indicate grouping, and p-values are shown within each panel.

Despite the urbanization-driven changes, seasons remained the main factor shaping the bacterial composition of the sponges. Samples collected during the dry season displayed a highly uniform bacterial community, regardless of the site’s exposure to urbanization pressures during the rainy season or host genera (Figure 2c). Under these conditions, the Burkholderiales order became the most abundant and evenly distributed taxon, accounting for over 50% in some samples (Figure 2a). Correlation analysis (Figure 2b) revealed that most taxa were negatively correlated with non-urbanized sites when considering seasonal influence, except Pseudomonadales, Bacillales (Figure 2d), and groups such as Flavobacteriales, Enterobacteriales, Aereomonadales, and Unclassified Gammaproteobacteria (Figure 2b), which were positively correlated and more abundant in non-urbanized areas. This suggests that seasons are significant forces shaping the sponge microbiome, even in more preserved environments, and parallels findings for other freshwater sponge species^58^. Urbanized sites showed few significant taxa changes across seasons, suggesting a similar microbial profile between urbanized sites, regardless of season, and that urbanization drives the sponge microbiome toward a taxonomic structure similar to that observed in the dry season (Figure S3). These findings suggest an uneven and dynamic microbiome in Amazonian sponges during the rainy season, particularly in non-urbanized regions. Under urbanized conditions and in dry-season samples, the microbiome remains quite similar, characterized by a high abundance of Burkholderiales (Figures 2a,d).

The sponge-associated eukaryotic organisms were more conserved across seasons (Figure 3a). During the rainy season, the main taxa were the sponge itself, unclassified eukaryotes, and metazoans. In the dry season, however, a fourth group was enriched: fungi, specifically Pezizomycotina through the classes Capnodiales and Dothideomycetes, which were the most differentially enriched (Figure 3b), along with other unclassified Pezizomycotina. These findings demonstrate that, unlike the bacterial community, the eukaryotic composition of freshwater sponges does not appear to be strongly affected by urbanization.

**Figure 3.**
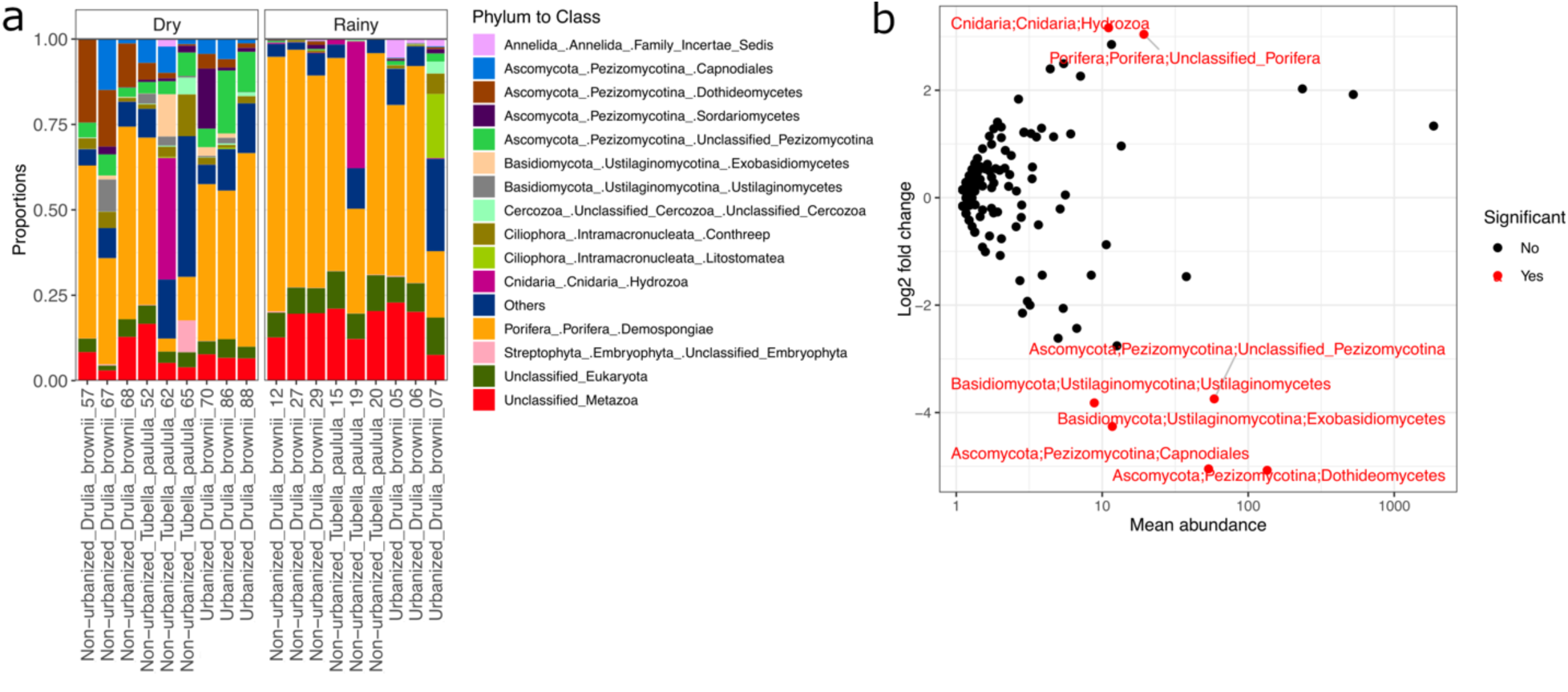
Eukaryotic community associated with Amazonian freshwater sponges across seasons. The figure (a) shows the taxonomical diversity of the 15 top eukaryotic classes in the sponge-associated consortia, shown as relative proportions; and (b) the differential abundance of the eukaryotes across seasons through a log10-scaled mean abundance (x-axis) and the log2 fold change (y-axis) of each taxon. Taxa significantly different in abundance between seasons (adjusted p < 0.05) are highlighted in red.

### Seasonal and urbanization-derived stress impact the gemmule microbiome: resilience or dysbiosis?

To understand how seasonality and urbanization could affect the long-term maintenance of the sponges, a metabarcoding sequencing of prokaryotic and eukaryotic gemmule-associated communities was conducted using 16S and 18S rRNA genes. The data indicated that the bacterial community associated to the gemmules was influenced by both seasonal changes and urbanization, especially the latter, contrasting with the whole microbiome, and these factors may interact to cause significant shifts in the presence and abundance of different taxa, depending on the stress levels they face (Figure 4a-b). During the rainy season, non-urbanized gemmules showed a high abundance of two main taxa: Unclassified Bacteria (53% on average) and Unclassified Gammaproteobacteria (29% on average), similar to what was found for *Ephydatia muelleri* unhatched gemmules ^59^. Together, these two groups accounted for over 75% of the bacterial composition across the three samples. Interestingly, the BLAST alignment of Unclassified Bacteria 16S against the NCBI database showed high identity to Erwiniaceae. This is followed by Enterobacteriales (3% on average) and Burkholderiales (2% on average) in smaller proportions (Figure 4a). Under urbanized conditions, the bacterial community in gemmules has undergone a significant shift, becoming richer and more even (Figure S4). Although Unclassified Gammaproteobacteria remained the dominant group, representing about 11% on average, Rhizobiales have become equally prevalent, accounting for roughly 10% of the bacteriome on average (Figure 4a). Comparing only species richness, without considering abundance, the non-urbanized sites harbor a more conserved microbiome (Figure 4c) than the urbanized sites (Figure 4d). It became evident when direct comparisons considering urbanization across seasons were made (Figure S4), which showed that the number of differentially abundant bacterial groups is far higher for urbanized sites (10-fold distance), indicating a clear instability of the consortia.

**Figure 4.**
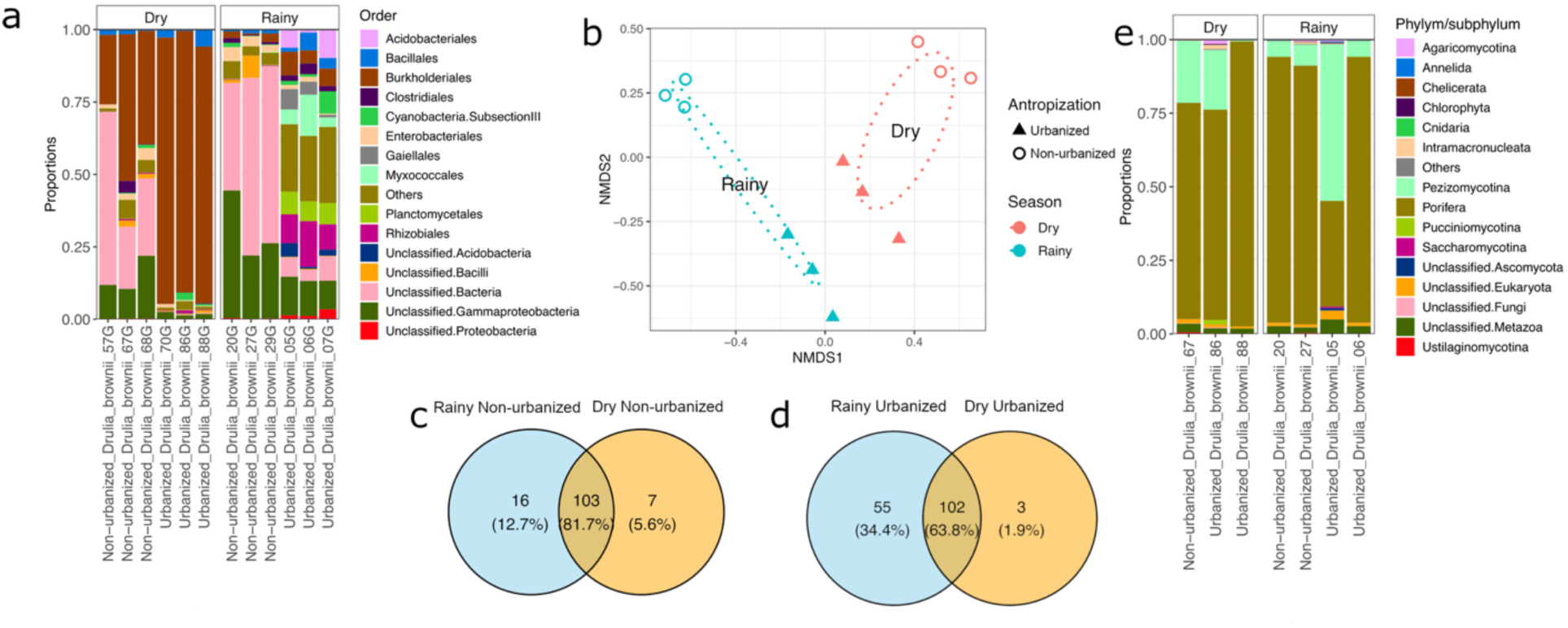
Metabarcoding of gemmule-associated consortia in freshwater sponges across seasons. Figure (a) shows the relative abundance (order level) of the 15 top prokaryotic organisms within the gemmules across both seasons for urbanized and non-urbanized environments; (b) NMDS of the prokaryotic composition of sponge gemmules based on Bray-Curtis dissimilarity, with points representing samples colored by season and shaped by site condition. Dotted ellipses denote 95% confidence intervals for seasonal groups; (c-d) Venn diagrams showing the shared organisms between the entire community and only non-urbanized sites across seasons; (e) relative abundance (phylum/subphylum level) of the 15 top eukaryotic organisms within the gemmules across both seasons.

During the dry season, gemmules collected from non-urbanized sites maintained the two main bacterial groups present in the rainy season, but in different proportions: Unclassified Bacteria averaged 35%, and Unclassified Gammaproteobacteria averaged 14%. Burkholderiales showed an increase in abundance, rising from about 2% in the rainy season to 37%. Despite some evidence already suggesting Burkholderiales as one of the dominant bacterial orders during gemmule hatching ^37^, extreme changes were observed in samples from urbanized sites during the dry season, where communities consisted almost completely of Burkholderiales (average 89%), with Bacillales contributing minimally (2.5%). These results indicate a stress-driven community shift, as the increase in Burkholderiales is suggested to be directly related to the level of stress imposed on the sponges. Specifically, urbanization has the least impact, while the dry season in urbanized samples exerts the greatest influence on the gemmules’ bacteriome. The gemmules-associated eukaryotic community, unlike the bacterial community, displayed high stability (Figure 4e).

### The Amazon sponge symbionts may be seasonally providing photoprotection, chemical defense, and enabling competition

To better understand the functional role of bacteria that compose the Amazon sponge microbiomes, we obtained MAGs from the metagenome. From the metagenomic contigs, 43 MAGs (Figure S5, Table S2) were retrieved, yielding 23 high-quality MAGs after species-level dereplication (ANI <95%). Additionally, five other cyanobacterial genomes were obtained from sponge tissue-directed cultivations using the same criteria (Table 1, Appendix S1). This inclusion was based in particular on the high number of photosynthesis genes in the dry-season whole-metagenome functional reconstruction for both sponges (Figure S6). The obtained MAGs were identified as belonging to 11 bacterial classes: Actinomycetales (7), Burkholderiales (5), Pseudomonadales (4), Bacillales (2), Flavobacteriales (1), Chitinophagales (1), Rhizobiales (1), Sphingomonadales (1), and Rickettsiales (1), and four other Nostocales and one Leptolyngbiales genomes were obtained from cultivated samples. MASH analysis revealed high genomic diversity for Actinomycetales and Burkholderiales, both of which are prevalent during the dry season; a considerable genomic distance between the Pseudomonadales MAGs, which belong to the Moraxellaceae and Pseudomonadaceae orders, and Flavobacteriales MAGs was observed. All the Burkholderiales were found to belong to the same genus, *Massilia*, while Actinomycetales was composed of six genera, and Pseudomonadales of two genera and one uncultured bacterium (Table 1).

Given the well-described role of bacterial symbionts in chemical defense within sponge consortia ^60,61^, the 28 MAGs were examined to identify the biosynthetic gene clusters (BGCs). At least 60 BGC types were identified across the MAGs, and despite the diversity, some MAGs exhibited similar biosynthetic profiles (Figure 5a, Supplementary File S1). Some BGCs, such as terpenes (95%), RiPPs and terpene precursors (73% each), and NRPS (65%), were highly distributed among MAGs (Figure 5b). The MAGs retrieved from the dry season exhibited greater BGC class diversity, with numerous NRPS, RiPPs, and terpenes (the richest class), suggesting an important biological role for them. MAGs from the rainy season, however, exhibited a lower biosynthetic richness and primarily focused on ecology-related BGC classes, such as terpenes, terpene precursors, N-siderophores, and aryl polyenes, with an increased representation of NRPS and other classes only in Pseudomonadales_20-0 (Figure S7).

**Figure 5.**
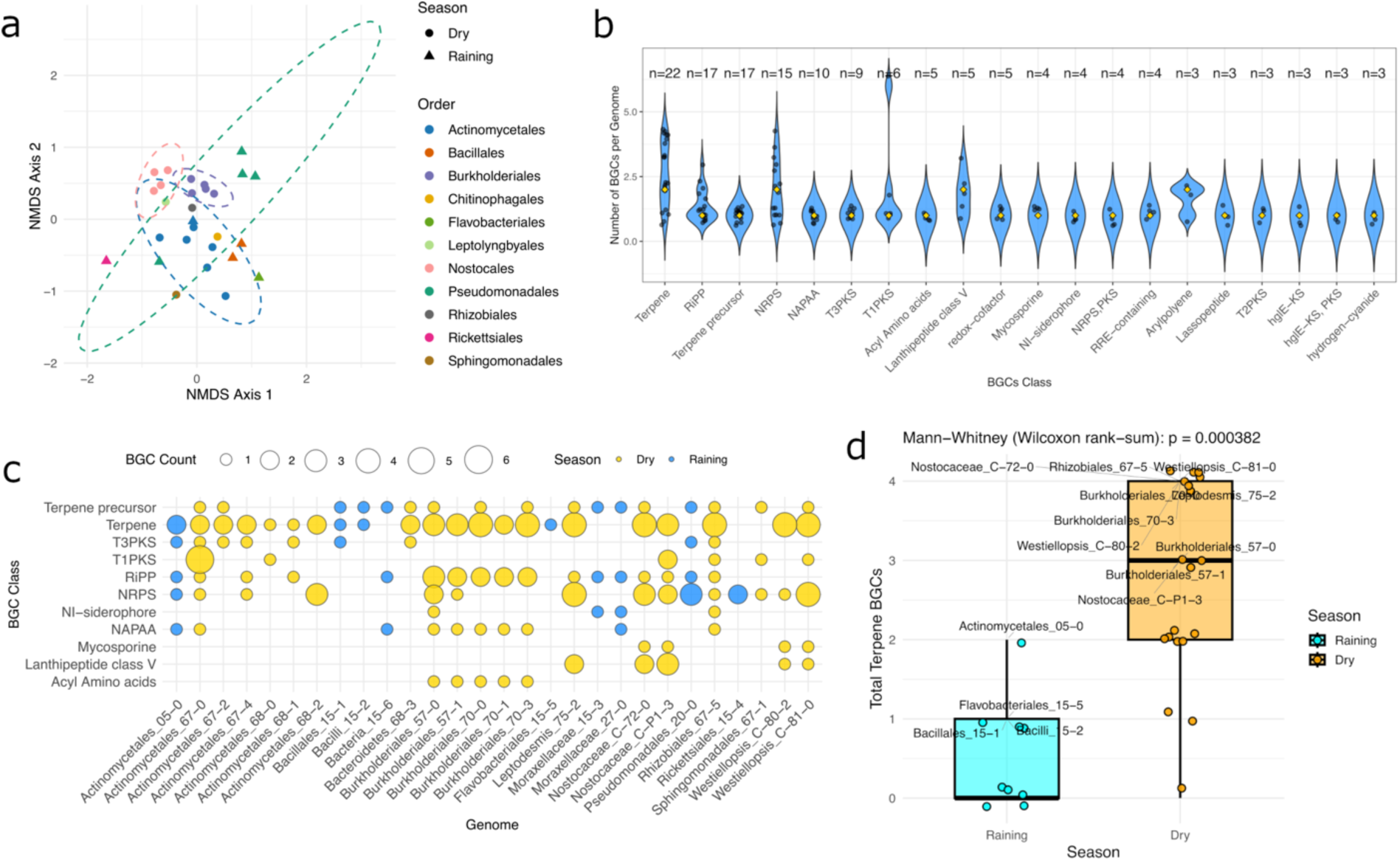
Diversity and distribution of BGCs across genomes and seasons. Figure (a) shows non-metric multidimensional scaling (NMDS) of BGC composition across genomes (Bray-Curtis on genome × BGC-class counts; k = 2). Points are genomes, colored by bacterial order and shaped by season. Dashed ellipses are 95% t-distribution confidence ellipses for each order (drawn only for groups with ≥ 3 genomes); (b) Violin plots show the distribution of counts for genomes containing each class (n = number of genomes). Black dots represent individual genome values, and yellow diamonds indicate medians; (c) bubble matrix of the top 11 BGC classes per genome. Rows are BGC classes and columns are genomes; bubble size scales with the number of BGCs detected in each genome-class cell. Color encodes season (Dry = yellow; Raining = blue). Empty cells indicate no BGCs of that class were found in that genome; (d) boxplots show the total terpene BGCs distribution for Raining (cyan) and Dry (orange); dots are individual genomes and labeled points mark genomes above the season-specific mean. The test used was the Wilcoxon rank-sum p-value.

The Burkholderiales MAGs, which were highly enriched during the dry season, shared conserved biosynthetic potential and clustered closely in the analysis (Figure 5a). They also represented the richest order in terpene BGCs (Figure 5c), followed by RiPPs, NAPPAs, and acyl amino acids. Similar evidence was observed for the cyanobacterial MAGs obtained from the dry season regarding terpene abundance, which also shared a conserved biosynthetic profile (Figure 5a) and were especially abundant in terpene BGCs (Figure 5c), NRPS, and lanthipeptide class v for the two Nostocaceae. Interestingly, most of the terpenes were found in MAGs from the dry season (Figure 5d), especially Burkholderiales and Nostocales, and all of them carry at least one carotenoid (Supplementary File S1). Given the wide distribution and abundance of terpenes and the conservative profile observed in dry-season MAGs, this evidence may suggest a specialized role for these compounds and a possible pressure for their maintenance over time in key bacteria.

Other bacterial classes, such as Actinomycetales, exhibited a wide dispersion of BGC profiles, and the same was observed for Pseudomonadales as a group (Figure 5a). Actinomycetales MAGs, with a dispersed and non-conserved profile (Figure 5a), showed a high diversity of BGCs, comprising the well-known bioactive-related classes NRPS, T1PKS, T3PKS, and RiPPs, as well as other ecology-related classes such as terpene and NAPAA BGCs. While some terpenes were shared among certain MAGs, such as the terpene isorenieratene, others were exclusive, such as the glycopeptidolipid identified in Actinomycetales_67-0. Two of the seven Actinomycetales MAGs (the 68-1 and 05-0, dry and rainy seasons) also possess hydrogen cyanide BGCs, such as those found for Pseudomonadales_20-0. Given the biosynthetic potential of Pseudomonadales_20-0 and its classification as a *Pseudomonas* strain, attempts were made to isolate the strain from fresh sponge tissue using Pseudomonas-specific cultivation methods (Appendix S1). However, none of the retrieved strains had sufficient 16S rRNA identity to be identified as the Pseudomonadales_20-0 strain.

### Microbe-microbe and microbe-environment systems mechanisms for host interaction and consortia supply

To shed light on the functional role of the MAGs in the consortia, a KEGG metabolic pathways reconstruction strategy was employed. The first analysis of the whole metagenomes functional annotation per condition showed an overall high number of genes related to the biosynthesis of secondary metabolites, microbial metabolism in diverse environments and two-component systems, highlighting the strong competitive, adaptive and regulatory microbial profile associated to sponges (Figure S6); curiously, as previously mentioned, a high number of photosynthesis genes were found in dry season for *D. brownii* and *T. paulula*, leading us to perform the cultivation of the cyanobacteria associated with sponge tissues (Appendix S2). A second functional analysis (Figure S8) showed clear distinctions between the MAGs: a first clade (A) harbor the four Pseudomonadales, five Burkholderiales and the Rhizobiales_67-5 MAGs, sub clustering them according to their respective seasons; a second clade (B) was found to harbor the seven Actinomycetales, two Bacillales and all other MAGs, not clustering them by season, but with most Actinomycetales in a separate subclade; a third clade (c), was cyanobacteria-specific, mainly influenced by photosynthesis genes. It was possible to find out that some important mechanisms for symbiotic associations, like Biofilm formation (Figure 6a), related to host colonization, and Two-component systems, a regulatory mechanism important for stress response (Figure 6a), were especially enriched for Burkholderiales and the MAG Pseudomonadales_20-0, both belonging to bacterial groups found to be highly abundant in sponges during dry and rainy seasons, respectively.

**Figure 6.**
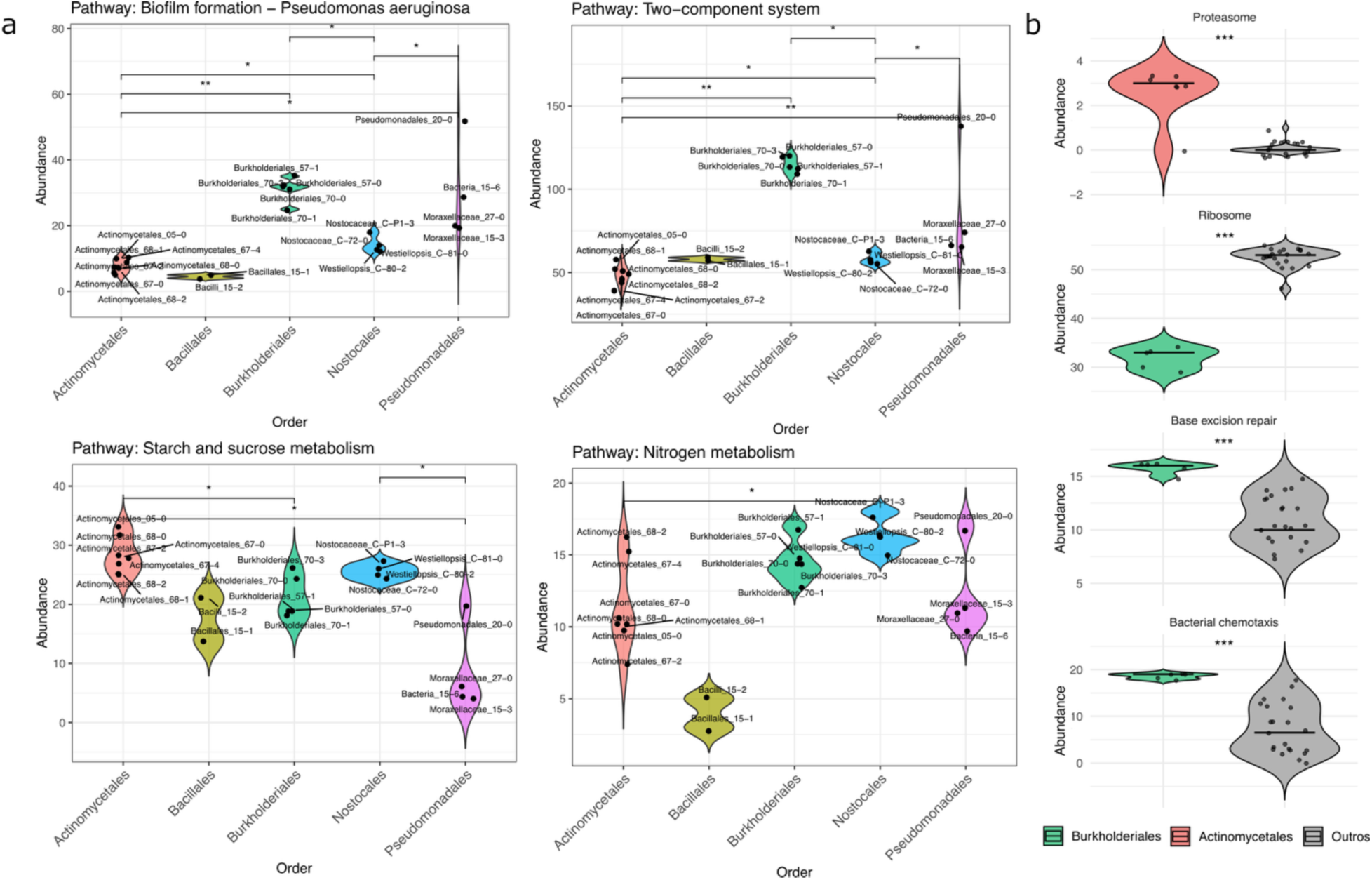
Overall and symbiosis-related KEGG metabolic pathways of the MAGs. Figure (a) shows violin plots with the distribution of abundance values for selected symbiosis-related pathways across bacterial orders with ≥2 genomes. Significance between orders was assessed by pairwise Wilcoxon rank-sum tests (p < 0.05), with asterisks indicating significance levels (p < 0.05 = *, p < 0.01 = **, p < 0.001 = ***); (b) comparison of selected KEGG pathway abundances between a test bacterial order against all other bacterial orders (“Others”). Each facet represents a pathway that showed a statistically significant difference (p < 0.05) in a Wilcoxon rank-sum test. Horizontal bars mark medians, and points represent individual genomes. Asterisks indicate significance levels (p < 0.05 = *, p < 0.01 = **, p < 0.001 = ***).

Burkholderiales MAGs also had significantly more genes related to Bacterial chemotaxis and Base excision repair (Figure 6b), which may be related to host colonization during the dry season and to DNA repair, given the stressful conditions during this season; interestingly, there was also a low number of ribosomal genes, supporting an oligotrophic profile (Figure 6b). The pathway Microbial metabolism in diverse environments was enriched for Actinomycetales MAGs and in small abundance for Bacillales MAGs (Figure S8), indicating an improved ability of Actinomycetales to cope with environmental challenges, as well as a lack of genomic specialization in Bacillales for this purpose. Actinomycetales also exhibited a considerable number of genes related to the Proteasome pathway, possibly as an adaptation to the oxidative stress resulting from prolonged exposure to UV light. The Carbon metabolism pathway (Figure S8), related to the ability to convert organic carbon sources into energy or its precursors, was found to be richer for Actinomycetales and Pseudomonadales_20-0 MAGs, suggesting a copiotroph-like role in the consortia. The Starch and sucrose metabolism pathway (Figure 6c) showed that most bacterial groups harbor a potentially strong ability to hydrolyze and metabolize complex carbohydrates into simple carbohydrates, even those with a small number of genes for the Carbon metabolism pathway, such as the two Bacillales MAGs, which was also found to be poorly involved in the Nitrogen metabolism (Figure 6a) and in the Vitamin B6 metabolism (Figure S8).

### The sponge holobiont may rely on key bacteria for the maintenance of the consortia’s core metabolism across seasons

The KEGG-based reconstruction of the complete metabolic modules, which considers only metabolic modules with all the necessary genes to work, showed a conserved profile mainly for two bacterial orders, Nostocales and Burkholderiales. Pseudomonadales, despite being dispersed, was found to overlap with Burkholderiales, which may have similar roles in different seasons (Figure 7a). When compared, dry season-related MAGs shared fewer complete modules (Figure 7b) than those of Pseudomonadales and Burkholderiales, obtained from different seasons (Figure 7c), supporting a possible shared role. A deeper analysis, examining order-specific pathways, revealed that the Burkholderiales-differentially abundant pathways were primarily associated with specific amino acids (lysine, tyrosine, and cysteine), especially glutamate, as well as vitamins and energy production (Figure 7d). In contrast, Actinomycetales exhibited a few, less specialized pathways, particularly those related to isoprenoid and pyridoxal phosphate biosynthesis (Figure 7e). Nostocales, as expected for cyanobacteria, were associated with nitrogen fixation and photosynthesis (Figure 7f). These pathways were among the top-enriched pathways between seasons, along with the biosynthesis of lysine and B6 vitamins in the dry season, as well as some amino acids (arginine, tyrosine, cysteine), heme, and alternative cytochrome oxidases (Figure 7g). Interestingly, alternative cytochrome oxidases were primarily found in Pseudomonadales and Burkholderiales MAGs, which also shared the Glyoxylate cycle module, as well as amino acid-related modules, particularly those related to glutamate and glutathione metabolism, biotin biosynthesis, and glutamate-based heme biosynthesis (Figure S9), a group of pathways involved in energy production.

**Figure 7.**
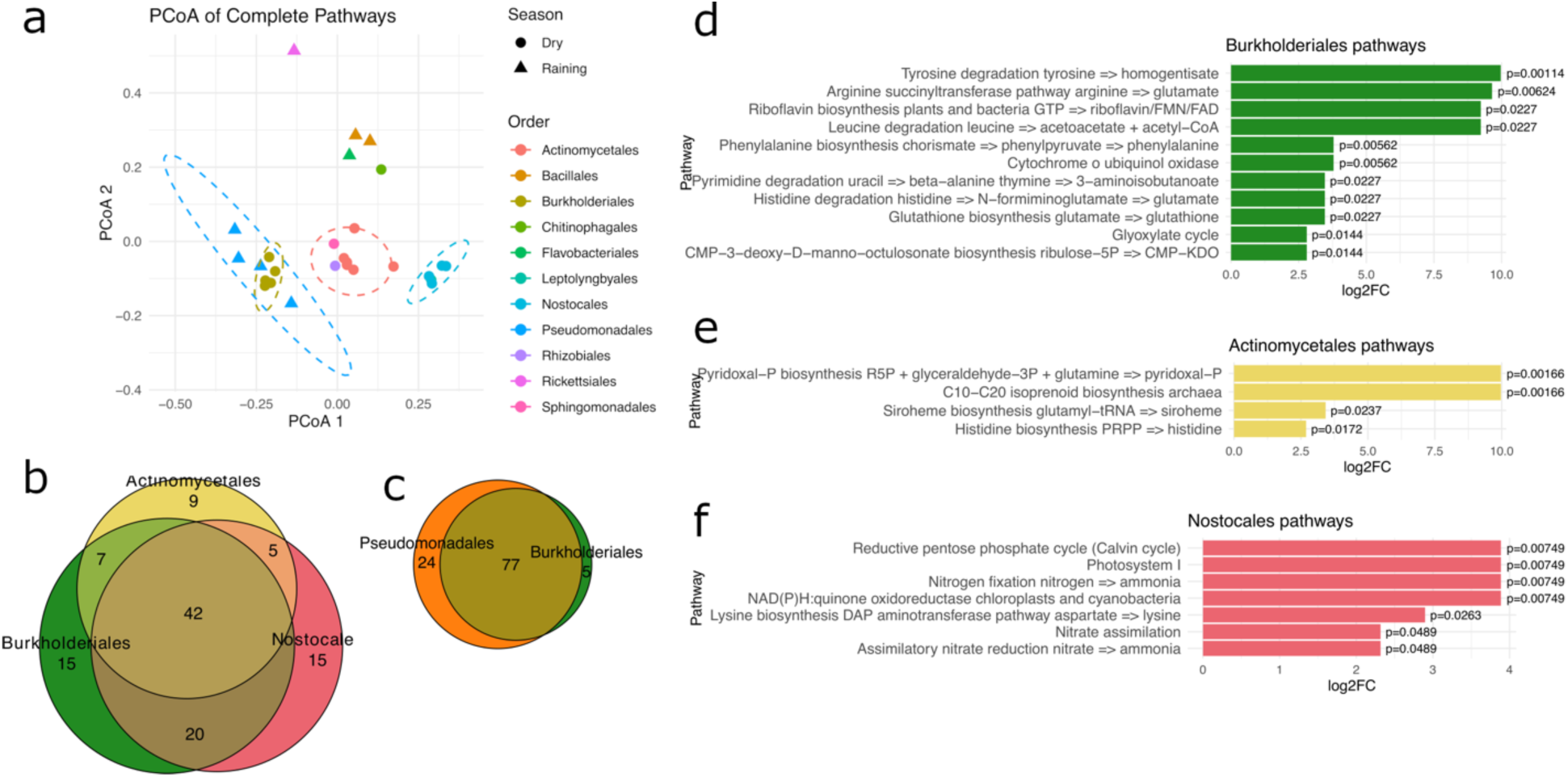
Complete overall and order-specific KEGG modules present in MAGs over seasons. Figure (a) shows PCoA based on Bray distances calculated from complete KEGG module profiles across genomes. Each point represents a genome, colored by bacterial order and shaped by season. Dashed ellipses are calculated at 95% confidence. Distances were computed on a presence/absence matrix. Statistical significance of compositional differences was tested using PERMANOVA (999 permutations); (b-c) Venn diagrams comparing pathways shared by the main orders; (d-f) Differentially enriched KEGG pathways in Actinomycetales, Nostocales, and Burkholderiales compared to other bacterial orders present in the dry season. Presence/absence complete KEGG modules were tested using Wilcoxon rank-sum tests, with Benjamini-Hochberg correction for multiple comparisons, and pathways with adjusted p-value < 0.05 and higher were retained. Bars represent the log2 fold change, and text labels show the adjusted p-values for each pathway.

### Symbiotic bacteria under carbon scarcity may have increased turnover capacity

Given the nutrient- and carbon-poor environment in which these sponges rely, and the scarcity scenario created by the dry season, a Carbohydrate-Active Enzymes (CAZymes) mining strategy was employed to gain insight into energy-related adaptations in sponge-associated bacteria. This approach revealed that MAGs from the dry season harbor a high number of carbohydrate-related enzymes (Figure 8a), with a mean abundance ranging from 37.5% to 87.5%. Burkholderiales were the richest bacterial group for CAZymes, followed by Cyanobacteria and one Bacteroidetes MAGs. In contrast, most of the MAGs from the rainy season showed a lower abundance, ranging from about 10% to 25%. Only two of these MAGs, namely Actinomycetales 05-5 and Flavobacteriales 15-5, showed higher abundances among season MAGs, between 37.5% and 50%. The overall composition profile observed separated four main clusters: three related to dry conditions and one related to the rainy condition (Figure 8b). Two additional isolated clusters are the Cyanobacteria cluster and the Burkholderiales cluster, which have opposite profiles, despite their richness in CAZymes. A third Actinomycetales cluster was shown to be more conserved, and the fourth one is composed of Pseudomonadales, Bacillales, and Rickettsiales. Curiously, Flavobacteriales was found to be a rainy season CAZymes-rich MAG and grouped among Burkholderiales.

**Figure 8.**
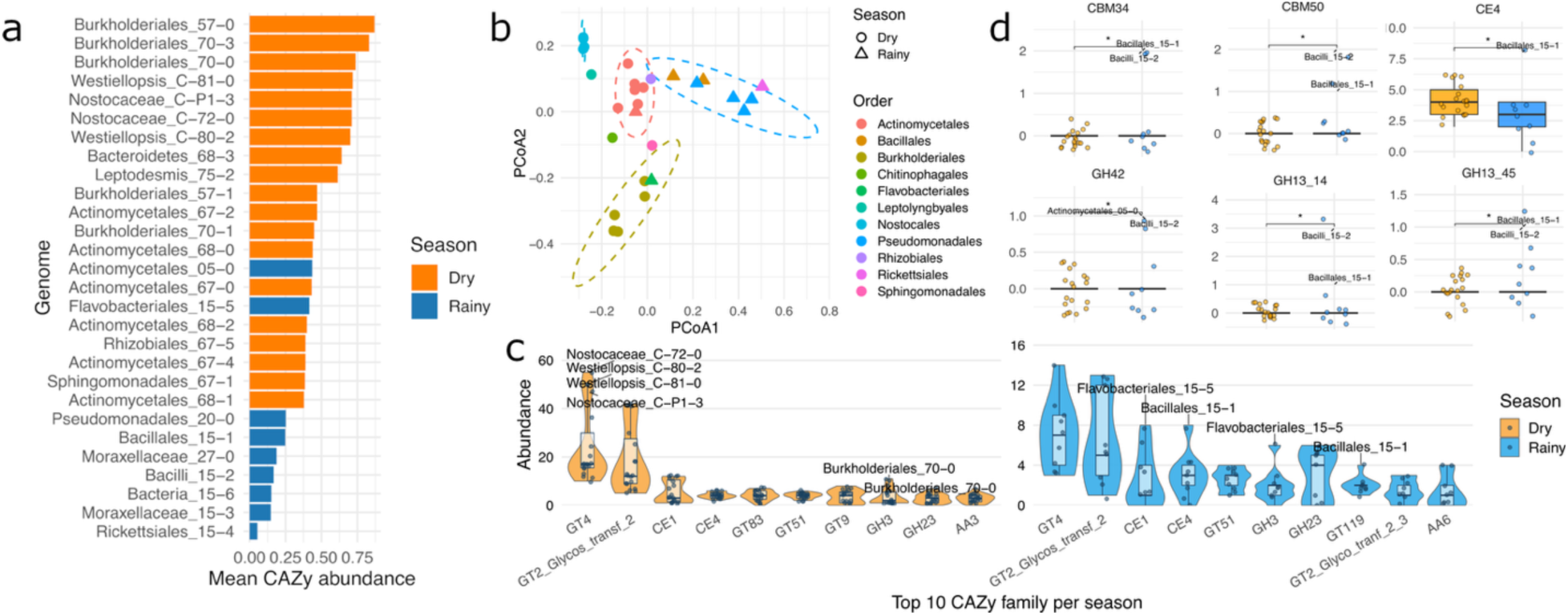
Seasonal variation in the abundance of CAZyme families across seasons. Figure (a) shows the average CAZy abundance per genome, colored by season, with genomes ordered by mean CAZy abundance and values calculated across all CAZy families; (b) PCoA based on Bray-Curtis distances of CAZyme family profiles across MAGs. Points represent genomes, colored by bacterial order and shaped by season. Dashed ellipses indicate 95% confidence intervals for orders with more than three genomes; (c) violin plots show the abundance distribution of the 10 most abundant CAZy per family across seasons, with boxplots indicating median and interquartiles. Points represent genomes, and labeled points mark high-abundance outliers (above 1× IQR from the upper quartile) identified per season and CAZy family; (d) boxplots show the distribution of CAZyme counts per genome for the seasons. Points represent genomes, and labeled points indicate outliers. Significance was assessed by a Wilcoxon rank-sum test (p ≤ 0.05), with asterisks denoting the significance.

The comparison of CAZymes distribution across seasons showed that the four richest families, comprising Glycosyltransferases (GT) and carbohydrate esterases (CE), remained stable over time (Figure 8c). Some, however, were found to be significantly increased during the dry season, a bias likely produced by Cyanobacteria MAGs due to their photosynthetic capacities. CE1 and CE4, in turn, are supposed to have a key role in both seasons, given their abundance and maintenance. Three families among the top 10 were replaced over the seasons: GT119, GT2_Glycos_transf_2_3, and an auxiliary activity AA6 are present among the wealthiest families in MAGs from the rainy season and absent from MAGs retrieved in the dry season, while AA3, GT9, and GT83 are the opposite. The other three families also remained over the seasons: two glycosyl hydrolases, GH3 and GH23, and a GT51. A more detailed analysis of comparing CAZymes present or absent over seasons showed Burkholderiales MAGs especially rich for CE1, GH3 and GT4; also, some enzymes were present only in the rainy season, specifically in Bacillales MAGs, like the carbohydrate binding modules CBM50 and CBM34, GH42, GH13_14 and GH13_45, and the aforementioned CE4 highly as enriched for the Bacillales_15-1, appointing the Bacillales’ role in carbohydrate management.

## DISCUSSION

This work, for the first time, conducted a taxonomic and functional characterization of the Amazon freshwater sponge microbiome across the two known seasons. Our metagenomic results showed that the naturally occurring microbiome in Amazon sponges is quite similar across the two studied species, despite their phylogenetic distance. In accordance with existing evidence, the Amazon sponge microbiomes were found to be heterogeneous even among intra-species individuals given the strong host-microbiome interactions ^26^, and were dominated by Pseudomonadales and Bacillales, bacterial groups that may play a crucial role in maintaining sponges. Flavobacteriales, present across the rainy season samples, especially in non-urbanized sites, are described as composing the microbiome of most of the freshwater sponges known to date ^37,38,59,62^. This microbiome profile, dominated by a few taxa, is commonly found in Low Microbial Abundance (LMA) sponges and is widespread across Demospongiae ^5^. Large-scale microbiome studies have shown that Gammaproteobacteria is the dominant class across marine sponges, commonly followed by Alphaproteobacteria ^7,26,63^. For freshwater sponges, a Gammaproteobacteria-rich microbiome was already described for the Spongillidae family through *Ephydatia muelleri* ^59^ and *Radiospongilla crateriformis*, *Eunapius fragilis*, and *Trochospongilla horrida* ^62^, while *Pseudomonas* strains were already isolated from freshwater sponges ^38,64^ and found to harbor a strong bioactivity in association with *Ephydatia fluviatilis* ^65^ and to be among the most frequently isolated genus in *T. varibilis* ^38^.

The results also suggest that urbanized sites influence the microbiome of Amazon freshwater sponges, producing a microbiome that is more even and more distant from that found at non-urbanized sites, consistent with similar evidence in Spongilla lacustris ^58^. Other studies with marine sponges in Patagonian Bay ^6^ and on the Atlantic coast of Brazil ^28^, both affected by wastewater, showed that chronic pollution produces changes in the microbiome through the reduction of the main bacterial groups, but these changes are commonly found in minor proportions. Evidence on three cosmopolitan freshwater sponge species exposed to pollutants showed that it also exhibited minor changes in their microbiome and enrichment of bacterial groups abundant in the surrounding water ^58,66,67^, however, a tropical freshwater sponge *Spongilla alba,* from Indian islands under two different degrees of potentially toxic elements, showed a more even microbiome and reduction of the dominant group under a more toxic environment ^68^. This evidence supports our findings and a dysbiosis-like event resulting from urbanization, but it’s unclear whether the microbiome changes are in fact a dysbiosis event or an acclimatization/adaptation in response to biotic stress, similar to what was found for marine sponges ^6^, especially considering the typical seasonal fluctuations in neotropical environments.

Amazon is known for its tropical two-season climate, characterized by a rainy-dry alternation over time ^69^. During the dry season, sponges are exposed to intense sunlight for many weeks, and the metagenomic analysis performed here found that seasonality is a significant force driving sponge microbiomes. The high temperatures and lack of access to water may lead sponge microbiomes to an even composition and, similar to what was found for the urbanization-derived stress, a depletion of the main bacterial groups. Similar findings for *Spongilla lacustris* support the seasonal pressure as the strongest force modulating freshwater sponge microbiomes ^58^. This pattern, however, does not apply to marine sponges, since seasonality does not significantly affect the microbiome in temperate climates, neither in European ^27,70,71^ nor in Asian sponges ^72^; some tropical sponges, however, undergo more expressive changes across seasonal pressures with temperature changing ^73^. Given that Amazon dry seasons do not produce high temperature fluctuations ^74^, instead producing extensive low water levels in the Amazon basin rivers, which create long-term stress driving the community to retain only well-adapted bacteria in the sponge microbiome.

The here-performed metagenomic analysis revealed that, during the dry season, in addition to the depletion of the main bacterial groups found in sponges, the microbiome undergoes a high enrichment of Burkholderiales across all the samples, independently of the urbanization degree or sponge species, already described as components of the marine sponges microbiome ^75,76^ and which may be related to the heat stress resistance ^77^. Five MAGs were taxonomically assigned to the genus *Massilia*, a recently described member of the Oxalobacteraceae (order Burkholderiales)^78^, which has been reported in extreme environments such as post-fire or fire-affected areas ^77,79,80^ where they quickly recover ^81,82^, desert soils ^83,84^, and exhibit high heat resistance. Despite being mainly characterized in heat-associated environments, these bacteria were also obtained from symbiotic-related conditions such as microbial mats from Antarctic samples in response to UV stress ^85^, associated with plants ^86,87^ and fungi ^88^, and were described as able to increase their abundance in a seasonally dependent way ^89^. It could explain the proliferation of these bacteria in such harsh conditions as the Amazon dry season, especially considering the symbiotic associations already demonstrated. Furthermore, Burkholderiales MAGs were found to be highly genomically diverse, and it’s unclear whether they were incorporated into the consortia as a group or diversified within the consortia. Bacterial symbiont genomes are known to be stable, as demonstrated for *Paraburkholderia* symbionts ^90^ and other organisms ^91–93^, however, intense stress exposure is also understood as a trigger for genomic diversification as an adaptive response ^94,95^, especially for oxidative stress ^96^, and it can help host resilience ^97^. This evidence on *Massilia* biology supports their association with Amazon freshwater sponges, which undergo annual extreme seasonal variations due to the Amazon’s fluctuating water levels and remain continually exposed to sunlight and UV stress during the Amazon dry season.

Freshwater sponges produce gemmules in response to harsh conditions, and these specialized structures also facilitate vertical transmission of microbes to the next generation ^37^. The microbiome of *D. brownii* gemmules was shown to be, under favorable conditions, dominated by unclassified bacteria and Gammaproteobacteria, as evidenced *Ephydatia muelleri* unhatched gemmules ^59^, and the unclassified bacteria were taxonomically closer to Erwiniaceae, a known group of symbionts ^98–100^. Urbanized sites, however, showed a strong influence on the microbial composition of these structures, leading to an even microbiome with reduced abundance of Unclassified bacteria and Gammaproteobacteria observed in non-urbanized sites. Interestingly, Burkholderiales abundance increased with the stress level the gemmules experienced, reaching extreme abundance in gemmule bacteriomes in dry-season urbanized-derived gemmules, similar to what was observed in the bulk microbiome. This suggests that although it’s not possible to observe an impact of urbanization on the bulk sponge microbiomes during the dry season, its actual influence is on the gemmule microbiomes. Despite being likely a sponge symbiont with a genomic content highly specialized for energy production and oxidative stress management, the over-enrichment of Burkholderiales may suggest a dysbiosis event resulting from exposure to two consecutive biotic and abiotic stressors, which could compromise the successful hatching of gemmules ^15^.

Evidence on *Spongilla lacustris* for gemmule hatching and development has shown that gemmules are dominated by a few bacterial orders, including Burkholderiales ^37^. The same study showed that the epibacterial community living on gemmule surfaces plays an important role in their maturation, and that the bacteriome diversifies during gemmule hatching. Tropical freshwater sponges have a faster but similar developmental process (in stages) when compared to temperate climate sponges (Calheira et al., 2019), however, to our knowledge, there are no studies evaluating the impact of dysbiosis-like events on the bulk microbiome of sponges during gemmule formation and what could be the long-term impact of it. The surrounding water has already been shown to affect the hatching of tropical freshwater sponges (Calheira et al., 2020), and urbanization has been correlated with the inability of gemmules to hatch in an Amazon Basin River (Volkmer-Ribeiro et al., 2008). Despite these findings providing evidence of possible dysbiosis in the gemmules of sponges living in urbanized sites, it is not possible to predict the negative impacts of this on gemmule hatching when environmental conditions become favorable or even in adult sponges in the long term.

The Firmicutes phylum, the second most abundant bacterial group in sponges during the rainy season, is primarily represented by the Bacillales order. This phylum, however, is not known for being widespread or abundant across marine sponge microbiomes ^26^. From the two Bacillales MAGs retrieved, the former, Bacillales_15-1, showed high 16S identity to the efficient feather-degrading *Ectobacillus* spJY-23, able to use keratin and other complex peptide molecules as a carbon source ^104^, and the second MAG, Bacillus_15-2, with *Exiguobacterium* sp. MH3, originally isolated from a rhizosphere ^105^, suggesting the sponge-associated strains possess a possible symbiotic role in the consortia and the ability to deal with complex molecules and fibrous, tough peptides ^106,107^. Bacillales MAGs, however, were not found to share core metabolism genes on a large scale, but especially for the pathway Starch and sucrose metabolism, Bacillus MAGs were shown to be at the same average as other bacterial groups, suggesting a streamlined core metabolism and an oligotroph-like metabolism, commonly found in Bacillus isolated from challenging oligotrophic environments ^108,109^. The Tapajós River, despite poor particulate and dissolved organic matter in the water, is suggested to be rich in algae and cyanobacteria ^110^, and to have C3 plants as the most important contributors to carbon sources in its lignin-rich waters ^111,112^. Given the close identity of one MAG to the feather-degrading *Ectobacillus* sp. JY-23, the oligotrophic-like profile and poor carbon-source profile of the Tapajós river, we hypothesized that Bacillales MAGs may be specialized in degrading complex organic matter, thereby contributing to carbon turnover in the consortia. Interestingly, one of the main sponge tissues, the spongin, is composed of collagen types I and III ^113^, and these bacteria may have a role in the breakdown and recycling of this tissue ^114^, actually acting as beneficial bacteria in the consortium, a hypothesis to be experimentally tested in future cultivation-based work.

To understand the potential role of sponge natural products, the obtained MAGs were analyzed, revealing a high richness of BGC classes, especially terpene and terpene precursor classes, which are well distributed and are described as abundant secondary metabolites in sponges ^115–118^. Terpenes were found across both seasons, but were richer in dry season MAGs, especially in Burkholderiales, and it was hypothesized that this abundance is mainly due to a possible photoprotective role ^117^, as known for carotenoids ^119–121^, able to protect organisms from oxidative stress ^122,123^ and useful in long-term UV exposure like the Amazon dry season for sponges. Terpenes also have ecological roles ^124,125^, being initially understood as related to symbiotic associations between invertebrates and their partners ^126^ and subsequently shown to modulate symbiotic microbial consortia ^127^, which could explain the larger abundance of this BGC class in the rainy season, when more intensive microbial interactions are expected.

The metabolic pathway reconstruction of the whole metagenome contigs also showed that, among the reconstructed pathways, the overall Microbial metabolism in diverse environments was the third most abundant, in accordance with the evidence on marine sponges, which shows the sponge microbiome as highly adaptable and resilient ^128,129^. Given that the main factor responsible for sponge adaptability is their associated microbes ^15^, the interaction between sponges and bacteria is a key element in adaptation. The data showed that, in Amazon sponges consortia, especially Actinomycetales, Burkholderiales and the Pseudomonadales harbor genes related to Microbial metabolism in diverse environments, while Burkholderiales and the Pseudomonadales_20-0 were found carrying many genes for Two-component systems, suggest a fine-tuned stimulus-response regulatory machinery ^130^ which is important for microbe-host interaction, such as Biofilm formation genes ^131^ and Chemotaxis ^132^, important mechanisms for host colonization and protection against stressful conditions ^133–137^, in addition to sharing many metabolic modules and being closer to each other, despite their different temporal abundance, than to other MAGs from the same season. Other pathways related to core metabolism, such as Nitrogen metabolism and Vitamin B6 metabolism, were identified as shared among Actinomycetales, Burkholderiales, the Pseudomonadales_20-0 MAG, and Cyanobacteria, providing insight into the possible role of cyanobacteria not as mutualists but as suppliers in the community, given its specialization in nitrogen-related pathways and photosynthesis, and highlighting the unique role of

Pseudomonadales_20-0 compared to other Pseudomonadales MAGs. Interestingly, all the bacteria in close association with sponges were found lacking CRISPR-Cas and without a high number of bacterial defense mechanisms (Figure S10), commonly found in higher abundance in environmental bacteria, given the need for competition ^138–140^ and in marine sponge-associated bacteria ^141–143^.

## CONCLUSIONS

This study brings the first evidence that the microbiome of freshwater sponges is strongly shaped by the Amazon environment, with two different sponge species sharing a similar microbiome despite their phylogenetic distance. It was also found that seasonal dynamics are a major force driving their microbiome across time, with stress-driven fluctuations emerging as an important factor modulating community composition and function. This seasonal imprint defines not only the adult sponge microbiome but also the bacterial consortia associated with gemmules, which may be the most vulnerable component of the holobiont. Under urbanized and dry-season conditions, gemmules exhibited marked reductions in diversity, indicating incipient dysbiosis that may compromise their role as reservoirs of microbial and functional continuity during environmental fluctuations. Among bacterial groups, Bacillales stood out as a distinctive feature of the Amazonian freshwater sponge microbiome. Their presence in adult sponges, but near absence in gemmules, suggests that they may play a specialized role linked to active host metabolism and to the degradation of complex organic matter, potentially contributing to nutrient turnover during favorable conditions. In contrast, Pseudomonadales and Burkholderiales may alternate in dominance across seasons: Pseudomonadales prevailed in rainy, non-urbanized environments, indicating a role in maintaining symbiotic balance under stable conditions, while Burkholderiales, a possible symbiont, expanded under dry-season and urbanized stress, suggesting an important but still poorly understood response to stress. Together, these patterns highlight how seasonality and disturbance may interact to restructure the sponge microbiome, with possible consequences for holobiont resilience over time. While our findings provide new insights into the ecological role of sponge-associated bacteria in Amazonian ecosystems, they also present limitations. Future efforts should aim to validate the genome-based functional predictions in vitro, explore the regulatory mechanisms driving host-symbiont interactions during seasonal transitions, and monitor the long-term impacts of urbanization on the persistence of these ancient holobionts.

## AUTHOR CONTRIBUTIONS

L.S.S. and M.P.C.S. conceptualized the study. J.L.A., G.S.T.F., D.A.G., and A.S. contributed to the sample and data obtention. L.S.S., J.L.A., and M.P.C.S. conducted the investigation. L.S.S. performed the formal analysis. A.S. and M.P.C.S. acquired funding. L.S.S. wrote the original draft, and J.L.A., G.S.T.F., R.A.B., D.A.G., A.S., and M.P.C.S. reviewed and included additional inputs to the manuscript.

## ACKNOWLEDGMENTS

We thank Jandria Gabriela Gusmão for her assistance with fieldwork and initial sample processing; Soraya Andrade and Silvanira Barbosa for their valuable support with all sequencing steps; Eduardo Lima for assistance with morphological procedures; and Bernardo Cintra for help with cyanobacterial cultivation. This research was supported by the brazilian Conselho Nacional de Desenvolvimento Científico e Tecnológico (CNPq) through the grant numbers 315214/2020-1 for M.P.C.S., 445350/2024-5 for Iwasa’i – Centro Avançado de Pesquisa e Inovação Biotecnológica da Amazônia Oriental, and 386039/2024-0 for L.S.S, and by Coordenação de Aperfeiçoamento de Pessoal de Nível Superior (CAPES) through fellow 88887.713111/2022-00 for L.S.S.

## DATA AVAILABILITY

The sequencing data produced in this work are available at the National Center for Biotechnology Information (https://www.ncbi.nlm.nih.gov/sra).

## REFERENCES

1. Yin, Z. et al. Sponge grade body fossil with cellular resolution dating 60 Myr before the Cambrian. Proc. Natl. Acad. Sci. U. S. A. 112, E1453–E1460 (2015).

2. Philippe, H. et al. Phylogenomics Revives Traditional Views on Deep Animal Relationships. Current Biology 19, 706–712 (2009).

3. Nielsen, C. Six major steps in animal evolution: are we derived sponge larvae? Evol. Dev. 10, 241–257 (2008).

4. de Voogd, N. J. et al. World Porifera Database. https://www.marinespecies.org/porifera (2025) doi:10.14284/359.

5. Busch, K. et al. Biodiversity, environmental drivers, and sustainability of the global deep-sea sponge microbiome. Nature Communications 2022 13:1 13, 1–16 (2022).

6. Gastaldi, M. et al. Holobiont dysbiosis or acclimatation? Shift in the microbial taxonomic diversity and functional composition of a cosmopolitan sponge subjected to chronic pollution in a Patagonian bay. PeerJ 12, e17707 (2024).

7. Cleary, D. F. R. et al. The sponge microbiome within the greater coral reef microbial metacommunity. Nature Communications 2019 10:1 10, 1–12 (2019).

8. Pronzato, R., Pisera, A. & Manconi. Fossil freshwater sponges: Taxonomy, geographic distribution, and critical review. Acta Palaeontol. Pol 62, 467–495 (2017).

9. Evans, K. L. & Montagnes, D. J. S. Freshwater sponge (Porifera: Spongillidae) distribution across a landscape: Environmental tolerances, habitats, and morphological variation. Invertebrate Biology 138, e12258 (2019).

10. van Soest, R. W. M. et al. Global Diversity of Sponges (Porifera). PLoS One 7, e35105 (2012).

11. Sugden, S., Holert, J., Cardenas, E., Mohn, W. W. & Stein, L. Y. Microbiome of the freshwater sponge Ephydatia muelleri shares compositional and functional similarities with those of marine sponges. ISME Journal 16, 2503–2512 (2022).

12. Wilkinson, C. R., Garrone, R. & Vacelet, J. Marine sponges discriminate between food bacteria and bacterial symbionts: electron microscope radioautography and in situ evidence. Proc. R. Soc. Lond. B Biol. Sci. 220, 519–528 (1984).

13. Müller, W. E. G., Wang, X. & Schröder, H. C. Paleoclimate and evolution: emergence of sponges during the neoproterozoic. Prog. Mol. Subcell. Biol. 47, 55–77 (2009).

14. Webster, N. S. & Thomas, T. The sponge hologenome. mBio 7, 135–151 (2016).

15. Pita, L., Rix, L., Slaby, B. M., Franke, A. & Hentschel, U. The sponge holobiont in a changing ocean: from microbes to ecosystems. Microbiome *2018* 6:1 6, 1–18 (2018).

16. Vacelet, J. & Donadey, C. Electron microscope study of the association between some sponges and bacteria. J. Exp. Mar. Biol. Ecol. 30, 301–314 (1977).

17. Tianero, M. D., Balaich, J. N. & Donia, M. S. Localized production of defence chemicals by intracellular symbionts of Haliclona sponges. Nature Microbiology 2019 4:7 4, 1149–1159 (2019).

18. Loh, T. L. & Pawlik, J. R. Chemical defenses and resource trade-offs structure sponge communities on Caribbean coral reefs. Proc. Natl. Acad. Sci. U. S. A. 111, 4151–4156 (2014).

19. Balskus, E. P. Sponge symbionts play defense. Nature Chemical Biology 2014 10:8 10, 611–612 (2014).

20. Flórez, L. V., Biedermann, P. H. W., Engl, T. & Kaltenpoth, M. Defensive symbioses of animals with prokaryotic and eukaryotic microorganisms. Nat. Prod. Rep. 32, 904–936 (2015).

21. De Goeij, J. M. et al. Surviving in a marine desert: The sponge loop retains resources within coral reefs. Science (1979). 342, 108–110 (2013).

22. Beazley, L. I., Kenchington, E. L., Murillo, F. J. & Sacau, M. D. M. Deep-sea sponge grounds enhance diversity and abundance of epibenthic megafauna in the Northwest Atlantic. ICES Journal of Marine Science 70, 1471–1490 (2013).

23. Bell, J. J. The functional roles of marine sponges. Estuar. Coast. Shelf Sci. 79, 341–353 (2008).

24. Engelberts, J. P. et al. Characterization of a sponge microbiome using an integrative genome-centric approach. ISME J. 14, 1100–1110 (2020).

25. Moitinho-Silva, L. et al. The sponge microbiome project. Gigascience 6, (2017).

26. Thomas, T. et al. Diversity, structure and convergent evolution of the global sponge microbiome. Nat. Commun. 7, 11870–11870 (2016).

27. Erwin, P. M., Coma, R., López-Sendino, P., Serrano, E. & Ribes, M. Stable symbionts across the HMA-LMA dichotomy: low seasonal and interannual variation in sponge-associated bacteria from taxonomically diverse hosts. FEMS Microbiol. Ecol. 91, (2015).

28. Hardoim, C. C. P., Hardoim, P. R., Lôbo-Hajdu, G., Custódio, M. R. & Thomas, T. The microbiome of the sponge Aplysina caissara in two sites with different levels of anthropogenic impact. FEMS Microbiol. Lett. 370, (2023).

29. Díez-Vives, C. et al. On the way to specificity - Microbiome reflects sponge genetic cluster primarily in highly structured populations. Mol. Ecol. 29, 4412–4427 (2020).

30. Griffiths, S. M. et al. Host genetics and geography influence microbiome composition in the sponge Ircinia campana. Journal of Animal Ecology 88, 1684–1695 (2019).

31. Gladkikh, A. S., Kalyuzhnaya, O. V., Belykh, O. I., Ahn, T. S. & Parfenova, V. V. Analysis of bacterial communities of two Lake Baikal endemic sponge species. Microbiology (Russian Federation) 83, 787–797 (2014).

32. Kulakova, N. V. et al. Brown Rot Syndrome and Changes in the Bacterial Сommunity of the Baikal Sponge Lubomirskia baicalensis. Microb. Ecol. 75, 1024–1034 (2018).

33. Parfenova, V. V. et al. Microbial community of freshwater sponges in Lake Baikal. Biology Bulletin 35, 374–379 (2008).

34. Belikov, S. et al. Diversity and shifts of the bacterial community associated with Baikal sponge mass mortalities. PLoS One 14, e0213926 (2019).

35. Gernert, C., Glöckner, F. O., Krohne, G. & Hentschel, U. Microbial diversity of the freshwater sponge Spongilla lacustris. Microb. Ecol. 50, 206–212 (2005).

36. Costa, R. et al. Evidence for Selective Bacterial Community Structuring in the Freshwater Sponge Ephydatia fluviatilis. Microb. Ecol. 65, 232–244 (2013).

37. Paix, B., van der Valk, E. & de Voogd, N. J. Dynamics, diversity, and roles of bacterial transmission modes during the first asexual life stages of the freshwater sponge Spongilla lacustris. *Environ*. Microbiome 19, 1–18 (2024).

38. Laport, M. S., Pinheiro, U. & Rachid, C. T. C. da C. Freshwater Sponge Tubella variabilis Presents Richer Microbiota Than Marine Sponge Species. Front. Microbiol. 10, (2019).

39. de Fernandes, M. G., Nascimento-Silva, G., Rozas, E. E., Hardoim, C. C. P. & Custódio, M. R. From Sea to Freshwater: Shared and Unique Microbial Traits in Sponge Associated Prokaryotic Communities. Curr. Microbiol. 82, 1–17 (2025).

40. Gonsior, M. et al. Chemodiversity of dissolved organic matter in the Amazon Basin. Biogeosciences 13, 4279–4290 (2016).

41. Tunno, I. et al. A morphological guide of neotropical freshwater sponge spicules for paleolimnological studies. Front. Ecol. Evol. 10, 1067432 (2023).

42. Volkmer-Ribeiro, C., Ezcurra de Drago, I., de Souza Machado, V. & Henry Sabaj, M. Drulia cristinae, new species of sponge from the rio Xingu, Amazonas Basin, Brazil (Porifera: Demospongiae: Poecilosclerida: Metaniidae Volkmer-Ribeiro, 1986). Proceedings of the Academy of Natural Sciences of Philadelphia 166, (2017).

43. Nunes, G., Custódio, M. R. & Pinheiro, U. A new species of Tubella (Porifera: Spongillidae) for the Brazilian Amazon: how misidentification can mask a potential species complex. Acta Amazon. 54, e54bc24187 (2024).

44. Folmer, O. et al. DNA primers for amplification of mitochondrial cytochrome c oxidase subunit I from diverse metazoan invertebrates. Mol. Mar. Biol. Biotechnol. (1994).

45. Katoh, K. & Standley, D. M. MAFFT Multiple Sequence Alignment Software Version 7: Improvements in Performance and Usability. Mol. Biol. Evol. 30, 772–780 (2013).

46. Hall, T. BioEdit version 7.2. 5. Ibis Biosciences, Carlsbad, CA, USA 10.1016/j.ifset.2004.06.001 (2013) doi:10.1016/j.ifset.2004.06.001.

47. Kumar, S., Stecher, G. & Tamura, K. MEGA7: Molecular Evolutionary Genetics Analysis Version 7.0 for Bigger Datasets. Mol. Biol. Evol. 33, 1870–1874 (2016).

48. De Coster, W., D’Hert, S., Schultz, D. T., Cruts, M. & Van Broeckhoven, C. NanoPack: visualizing and processing long-read sequencing data. Bioinformatics 34, 2666–2669 (2018).

49. Bengtsson-Palme, J. et al. metaxa2: improved identification and taxonomic classification of small and large subunit rRNA in metagenomic data. Mol. Ecol. Resour. 15, 1403–1414 (2015).

50. Kolmogorov, M., Yuan, J., Lin, Y. & Pevzner, P. A. Assembly of long, error-prone reads using repeat graphs. Nature Biotechnology 2019 37:5 37, 540–546 (2019).

51. Li, H. Minimap2: pairwise alignment for nucleotide sequences. Bioinformatics 34, 3094–3100 (2018).

52. Pan, S., Zhao, X. M. & Coelho, L. P. SemiBin2: self-supervised contrastive learning leads to better MAGs for short- and long-read sequencing. Bioinformatics 39, i21–i29 (2023).

53. Parks, D. H., Imelfort, M., Skennerton, C. T., Hugenholtz, P. & Tyson, G. W. CheckM: assessing the quality of microbial genomes recovered from isolates, single cells, and metagenomes. Genome Res. 25, 1043–1055 (2015).

54. Olm, M. R., Brown, C. T., Brooks, B. & Banfield, J. F. dRep: a tool for fast and accurate genomic comparisons that enables improved genome recovery from metagenomes through de-replication. ISME J. 11, 2864–2868 (2017).

55. Cantalapiedra, C. P., Hern̗andez-Plaza, A., Letunic, I., Bork, P. & Huerta-Cepas, J. eggNOG-mapper v2: Functional Annotation, Orthology Assignments, and Domain Prediction at the Metagenomic Scale. Mol. Biol. Evol. 38, 5825–5829 (2021).

56. Blin, K. et al. antiSMASH 7.0: new and improved predictions for detection, regulation, chemical structures and visualisation. Nucleic Acids Res. 51, W46–W50 (2023).

57. Tesson, F. et al. Systematic and quantitative view of the antiviral arsenal of prokaryotes. Nature Communications 2022 13:1 13, 1–10 (2022).

58. Lhomme, C., Paix, B. & de Voogd, N. J. Host senescence and urbanized conditions drive bacterial communities of the freshwater sponge Spongilla lacustris. bioRxiv 10.64898/2026.01.07.698122 (2026) doi:10.64898/2026.01.07.698122.

59. Kenny, N. J. et al. Tracing animal genomic evolution with the chromosomal-level assembly of the freshwater sponge Ephydatia muelleri. Nature Communications 2020 11:1 11, 1–11 (2020).

60. Rust, M. et al. A multiproducer microbiome generates chemical diversity in the marine sponge Mycale hentscheli. Proc. Natl. Acad. Sci. U. S. A. 117, 9508–9518 (2020).

61. Zhang, S. et al. Comparative metabolomic analysis reveals shared and unique chemical interactions in sponge holobionts. Microbiome 10, (2022).

62. Keleher, J. G. et al. Freshwater sponges in the southeastern U.S. harbor unique microbiomes that are influenced by host and environmental factors. PeerJ 13, e18807 (2025).

63. Turon, M. et al. Microbiome changes through the ontogeny of the marine sponge Crambe crambe. *Environ*. Microbiome 19, 1–17 (2024).

64. Clark, C. M. et al. Relationship between bacterial phylotype and specialized metabolite production in the culturable microbiome of two freshwater sponges. ISME Communications 2, 22 (2022).

65. Keller-Costa, T., Jousset, A., Van Overbeek, L., Van Elsas, J. D. & Costa, R. The Freshwater Sponge Ephydatia fluviatilis Harbours Diverse Pseudomonas Species (Gammaproteobacteria, Pseudomonadales) with Broad-Spectrum Antimicrobial Activity. PLoS One 9, e88429 (2014).

66. Rizzo, C. et al. Microbiome and pollutants in the freshwater sponges Ephydatia muelleri (Lieberkühn, 1856) and Spongilla lacustris (Linnaeus, 1758) from the sub-Arctic Pasvik river (Northern Fennoscandia). Environ. Res. 273, 121126 (2025).

67. Cartwright, A., Dooley, J. S. G., McGonigle, C. D. & Arnscheidt, J. How suitable is freshwater sponge Ephydatia fluviatilis (Linnaeus, 1759) for time-integrated biomonitoring of microbial water quality? Access Microbiol. 6, 000691.v4 (2024).

68. Bhattacharya, D. et al. Bacterial diversity in the freshwater sponges of Sundarban and their potential role in biomonitoring toxic element pollution. Microbiol. Spectr. 10.1128/SPECTRUM.02149-25 (2025) doi:10.1128/SPECTRUM.02149-25.

69. Correa, S. B. et al. Biotic Indicators for Ecological State Change in Amazonian Floodplains. Bioscience 72, 753 (2022).

70. Lamb, C. E. & Watts, J. E. M. Microbiome species diversity and seasonal stability of two temperate marine sponges Hymeniacidon perlevis and Suberites massa. *Environ*. Microbiome 18, (2023).

71. Erwin, P. M., Pita, L., López-Legentil, S. & Turon, X. Stability of sponge-associated bacteria over large seasonal shifts in temperature and irradiance. Appl. Environ. Microbiol. 78, 7358–7368 (2012).

72. Gan, B. et al. Dynamic microbiome diversity shaping the adaptation of sponge holobionts in coastal waters. Microbiol. Spectr. 12, (2024).

73. Hayami, Y., Ambalavanan, L., Zainathan, S. C., Danish-Daniel, M. & Iehata, S. Temporal dynamics of the bacterial community structure and functions associated with marine sponges collected off Karah Island, Terengganu, Malaysia. The Microbe 5, 100202 (2024).

74. Marengo, J. A. et al. Changes in Climate and Land Use Over the Amazon Region: Current and Future Variability and Trends. Front. Earth Sci. (Lausanne). 6, 425317 (2018).

75. Hardoim, C. C. P. et al. Diversity of bacteria in the marine sponge aplysina fulva in brazilian coastal waters. Appl. Environ. Microbiol. 75, 3331–3343 (2009).

76. Trindade-Silva, A. E. et al. Taxonomic and Functional Microbial Signatures of the Endemic Marine Sponge Arenosclera brasiliensis. PLoS One 7, e39905 (2012).

77. De Castro-Fernández, P. et al. How does heat stress affect sponge microbiomes? Structure and resilience of microbial communities of marine sponges from different habitats. Front. Mar. Sci. 9, (2023).

78. La Scola, B., Birtles, R. J., Mallet, M. N. & Raoult, D. Massilia timonae gen. nov., sp. nov., isolated from blood of an immunocompromised patient with cerebellar lesions. J. Clin. Microbiol. 36, 2847–2852 (1998).

79. Nelson, A. R. et al. Wildfire impact on soil microbiome life history traits and roles in ecosystem carbon cycling. ISME Communications 4, 108 (2024).

80. Whitman, T. et al. Soil bacterial and fungal response to wildfires in the Canadian boreal forest across a burn severity gradient. Soil Biol. Biochem. 138, 107571 (2019).

81. Pulido-Chavez, M. F. et al. Rapid bacterial and fungal successional dynamics in first year after chaparral wildfire. Mol. Ecol. 32, 1685–1707 (2023).

82. Soria, R. et al. Short-Term Response of Soil Bacterial Communities after Prescribed Fires in Semi-Arid Mediterranean Forests. Fire 2023, Vol. 6, Page 145 6, 145 (2023).

83. Manni, A., Filali-Maltouf, A., Manni, A. & Filali-Maltouf, A. Diversity and bioprospecting for industrial hydrolytic enzymes of microbial communities isolated from deserted areas of south-east Morocco. AIMS Microbiology 2022 1:5 8, 5–25 (2022).

84. Selmani, Z. et al. Culturing the desert microbiota. Front. Microbiol. 14, 1098150 (2023).

85. Shaffer, J. M. C., Giddings, L. A., Samples, R. M. & Mikucki, J. A. Genomic and phenotypic characterization of a red-pigmented strain of Massilia frigida isolated from an Antarctic microbial mat. Front. Microbiol. 14, 1156033 (2023).

86. Liu, X. L. et al. A primary assessment of the endophytic bacterial community in a xerophilous moss (Grimmia montana) using molecular method and cultivated isolates. Brazilian Journal of Microbiology 45, 165–173 (2014).

87. Ofek, M., Hadar, Y. & Minz, D. Ecology of Root Colonizing Massilia (Oxalobacteraceae). PLoS One 7, e40117 (2012).

88. Scheublin, T. R., Sanders, I. R., Keel, C. & Van Der Meer, J. R. Characterisation of microbial communities colonising the hyphal surfaces of arbuscular mycorrhizal fungi. ISME J. 4, 752–763 (2010).

89. Xu, A., Liu, C., Zhao, S., Song, Z. & Sun, H. Dynamic distribution of Massilia spp. in sewage, substrate, plant rhizosphere/phyllosphere and air of constructed wetland ecosystem. Front. Microbiol. 14, 1211649 (2023).

90. Noh, S. et al. Reduced and Nonreduced Genomes in Paraburkholderia Symbionts of Social Amoebas. mSystems 7, (2022).

91. Clayton, A. L., Jackson, D. G., Weiss, R. B. & Dale, C. Adaptation by Deletogenic Replication Slippage in a Nascent Symbiont. Mol. Biol. Evol. 33, 1957–1966 (2016).

92. Otero-Bravo, A. & Sabree, Z. L. Multiple concurrent and convergent stages of genome reduction in bacterial symbionts across a stink bug family. Scientific Reports 2021 11:1 11, 1–15 (2021).

93. Hansen, A. K., Percy, D. M., Miao, S. & Degnan, P. H. Effect of Oceanic Islands on an Insect Symbiont Genome in Transition to a Host-Restricted Lifestyle. Genome Biol. Evol. 17, (2025).

94. Hoffmann, A. A. & Hercus, M. J. Environmental Stress as an Evolutionary Force. Bioscience 50, 217–226 (2000).

95. Ament-Velásquez, S. L. et al. The Dynamics of Adaptation to Stress from Standing Genetic Variation and de novo Mutations. Mol. Biol. Evol. 39, (2022).

96. Boles, B. R. & Singh, P. K. Endogenous oxidative stress produces diversity and adaptability in biofilm communities. Proc. Natl. Acad. Sci. U. S. A. 105, 12503–12508 (2008).

97. Tougeron, K. & Iltis, C. Impact of heat stress on the fitness outcomes of symbiotic infection in aphids: a meta-analysis. Proceedings of the Royal Society B: Biological Sciences 289, 20212660 (2022).

98. Siden-Kiamos, I. et al. Dynamic interactions between the symbiont Candidatus Erwinia dacicola and its olive fruit fly host Bactrocera oleae. Insect Biochem. Mol. Biol. 146, (2022).

99. Nia, T. & Id, N. Olive fruit fly and its obligate symbiont Candidatus Erwinia dacicola: Two new symbiont haplotypes in the Mediterranean basin. PLoS One 16, e0256284 (2021).

100. Estes, A. M., Hearn, D. J., Agrawal, S., Pierson, E. A. & Dunning Hotopp, J. C. Comparative genomics of the Erwinia and Enterobacter olive fly endosymbionts. Scientific Reports 2018 8:1 8, 1–13 (2018).

101. Calheira, L., Lanna, E. & Pinheiro, U. Tropical freshwater sponges develop from gemmules faster than their temperate-region counterparts. Zoomorphology 138, 425–436 (2019).

102. Calheira, L., Santos, P. J. P. & Pinheiro, U. Hatchability of gemmules of two Neotropical freshwater sponges (Porifera: Spongillida). Iheringia Ser. Zool. 110, e2020001 (2020).

103. Volkmer-Ribeiro, C., Batista, T. C. A., Melão, M. G. G. & Fonseca-Gessner, A. A. Anthropically dislodged assemblages of sponges (Porifera: Demospongiae) in the river Araguaia at Araguatins, Tocantins State, Brazil. Acta Limnologica Brasiliensia 20, 169–175 (2008).

104. Peng, S. et al. Isolation of a novel feather-degrading Ectobacillus sp. JY-23 strain and characterization of a new keratinase in the M4 metalloprotease family. Microbiol. Res. 274, (2023).

105. Tang, J., Zhang, Y., Meng, H., Xue, Z. & Ma, J. Complete Genome Sequence of Exiguobacterium sp. Strain MH3, Isolated from Rhizosphere of Lemna minor. Genome Announc. 1, 1059–1072 (2013).

106. Shao, X. et al. Culture Condition Optimization and Pilot Scale Production of the M12 Metalloprotease Myroilysin Produced by the Deep-Sea Bacterium Myroides profundi D25. Molecules 2015, Vol. 20, Pages 11891-11901 20, 11891–11901 (2015).

107. Chen, X. L. et al. Ecological function of myroilysin, a novel bacterial M12 metalloprotease with elastinolytic activity and a synergistic role in collagen hydrolysis, in biodegradation of deep-sea high-molecular-weight organic nitrogen. Appl. Environ. Microbiol. 75, 1838–1844 (2009).

108. Alcaraz, L. D. et al. The genome of Bacillus coahuilensis reveals adaptations essential for survival in the relic of an ancient marine environment. Proc. Natl. Acad. Sci. U. S. A. 105, 5803–5808 (2008).

109. Alcaraz, L. D. et al. Understanding the evolutionary relationships and major traits of Bacillus through comparative genomics. BMC Genomics 11, 332 (2010).

110. Li, S. et al. Distinct Non-conservative Behavior of Dissolved Organic Matter after Mixing Solimões/Negro and Amazon/Tapajós River Waters. ACS ES and T Water 3, 2083–2095 (2023).

111. Bertassoli, D. J. et al. The fate of carbon in sediments of the Xingu and Tapajós clearwater rivers, eastern Amazon. Front. Mar. Sci. 4, 241456 (2017).

112. Farella, N., Lucotte, M., Louchouarn, P. & Roulet, M. Deforestation modifying terrestrial organic transport in the Rio Tapajós, Brazilian Amazon. Org. Geochem. 1443–1458 (2001).

113. Ehrlich, H. et al. Discovery of mammalian collagens I and III within ancient poriferan biopolymer spongin. Nature Communications 2025 16:1 16, 1–13 (2025).

114. Esteves, A. I. S., Cullen, A. & Thomas, T. Competitive interactions between sponge-associated bacteria. FEMS Microbiol. Ecol. 93, 8 (2017).

115. Xu, J. et al. Terpenoids from the Sponge Sarcotragus sp. Collected in the South China Sea. J. Nat. Prod. 86, 330–339 (2023).

116. Dyshlovoy, S. A. et al. New diterpenes from the marine sponge Spongionella sp. overcome drug resistance in prostate cancer by inhibition of P-glycoprotein. Scientific Reports 2022 12:1 12, 1–13 (2022).

117. Martignago, C. C. S. et al. Terpenes extracted from marine sponges with antioxidant activity: a systematic review. Nat. Prod. Bioprospect. 13, 23 (2023).

118. Yuan, Y., Lei, Y., Xu, M., Zhao, B. & Xu, S. Bioactive Terpenes from Marine Sponges and Their Associated Organisms. Mar. Drugs 23, 96 (2025).

119. Paredes Contreras, B. V. et al. Enhanced UV-B photoprotection activity of carotenoids from the novel Arthrobacter sp. strain LAPM80 isolated from King George Island, Antarctica. Heliyon 11, e41400 (2025).

120. Reis-Mansur, M. C. P. P. et al. Carotenoids from UV-resistant Antarctic Microbacterium sp. LEMMJ01. Scientific Reports 2019 9:1 9, 1–14 (2019).

121. Liu, H. et al. UV-B irradiation differentially regulates terpene synthases and terpene content of peach. Plant Cell Environ. 40, 2261–2275 (2017).

122. Singh, B. K., Tripathi, M., Chaudhari, B. P., Pandey, P. K. & Kakkar, P. Natural Terpenes Prevent Mitochondrial Dysfunction, Oxidative Stress and Release of Apoptotic Proteins during Nimesulide-Hepatotoxicity in Rats. PLoS One 7, e34200 (2012).

123. Ayala-Ruiz, L. A. et al. Role of the major terpenes of Callistemon citrinus against the oxidative stress during a hypercaloric diet in rats. Biomedicine & Pharmacotherapy 153, 113505 (2022).

124. Kandasamy, D. et al. Conifer-killing bark beetles locate fungal symbionts by detecting volatile fungal metabolites of host tree resin monoterpenes. PLoS Biol. 21, e3001887 (2023).

125. Ivanisevic, J. et al. Biochemical Trade-Offs: Evidence for Ecologically Linked Secondary Metabolism of the Sponge Oscarella balibaloi. PLoS One 6, e28059 (2011).

126. Kokke, W. C. et al. On the origin of terpenes in symbiotic associations between marine invertebrates and algae (zooxanthellae). Culture studies and an application of 13C/12C isotope ratio mass spectrometry. Journal of Biological Chemistry 259, 8168–8173 (1984).

127. Ali, M. et al. Overexpression of Terpenoid Biosynthesis Genes From Garden Sage (Salvia officinalis) Modulates Rhizobia Interaction and Nodulation in Soybean. Front. Plant Sci. 12, 783269 (2021).

128. Botté, E. S. et al. Changes in the metabolic potential of the sponge microbiome under ocean acidification. Nat. Commun. 10, 4134 (2019).

129. Chai, G., Li, J. & Li, Z. The interactive effects of ocean acidification and warming on bioeroding sponge Spheciospongia vesparium microbiome indicated by metatranscriptomics. Microbiol. Res. 278, 127542 (2024).

130. Groisman, E. A. Feedback Control of Two-Component Regulatory Systems. Annu. Rev. Microbiol. 70, 103 (2016).

131. Liu, C., Sun, D., Zhu, J. & Liu, W. Two-component signal transduction systems: A major strategy for connecting input stimuli to biofilm formation. Front. Microbiol. 10, 420518 (2019).

132. Kirby, J. R. Chemotaxis-like regulatory systems: unique roles in diverse bacteria. Annu. Rev. Microbiol. 63, 45–59 (2009).

133. Yang, L. et al. Mechanisms of rhizosphere plant-microbe interactions: molecular insights into microbial colonization. Front. Plant Sci. 15, 1491495 (2024).

134. Flemming, H. C. et al. The biofilm matrix: multitasking in a shared space. Nat. Rev. Microbiol. 21, 70–86 (2023).

135. Chagas, F. O., Pessotti, R. D. C., Caraballo-Rodríguez, A. M. & Pupo, M. T. Chemical signaling involved in plant-microbe interactions. Chem. Soc. Rev. 47, 1652–1704 (2018).

136. Allard-Massicotte, R. et al. Bacillus subtilis early colonization of Arabidopsis thaliana roots involves multiple chemotaxis receptors. mBio 7, (2016).

137. Arnaouteli, S., Bamford, N. C., Stanley-Wall, N. R. & Kovács, Á. T. Bacillus subtilis biofilm formation and social interactions. Nat. Rev. Microbiol. 19, 600–614 (2021).

138. Beavogui, A. et al. The defensome of complex bacterial communities. Nature Communications 2024 15:1 15, 1–15 (2024).

139. Xiao, W., Weissman, J. L. & Johnson, P. L. F. Ecological drivers of CRISPR immune systems. mSystems 9, (2024).

140. Smith, W. P. J., Wucher, B. R., Nadell, C. D. & Foster, K. R. Bacterial defences: mechanisms, evolution and antimicrobial resistance. Nature Reviews Microbiology 2023 21:8 21, 519–534 (2023).

141. Wang, W. et al. Decoupling of strain- and intrastrain-level interactions of microbiomes in a sponge holobiont. Nature Communications 2024 15:1 15, 1–17 (2024).

142. Robbins, S. J. et al. A genomic view of the microbiome of coral reef demosponges. ISME J. 15, 1641 (2021).

143. Horn, H. et al. An Enrichment of CRISPR and other defense-related features in marine sponge-associated microbial metagenomes. Front. Microbiol. 7, 1751 (2016).

