## Supporting information for "Two Amazon freshwater sponges, *Drulia brownii* and *Tubella paulula*, share a resilient microbiome strongly shaped by seasonality and urbanization"

L. S. S.[1], J. L. A.[1,2], G. S. T. F.[3], R. A. B.[4], D. A. G.[4], A. S.[1,4], M. P. C. S.[1,4]

[1]Institute of Biological Sciences, Universidade Federal do Pará.

[2] Instituto Federal do Pará, Belém, PA, Brazil.

[3] Institute of Science and Technology of Waters, Universidade Federal do Oeste do Pará, Santarém, PA, Brazil.

[4] Parque de Ciência e Tecnologia Guamá, Belém, PA, Brazil.

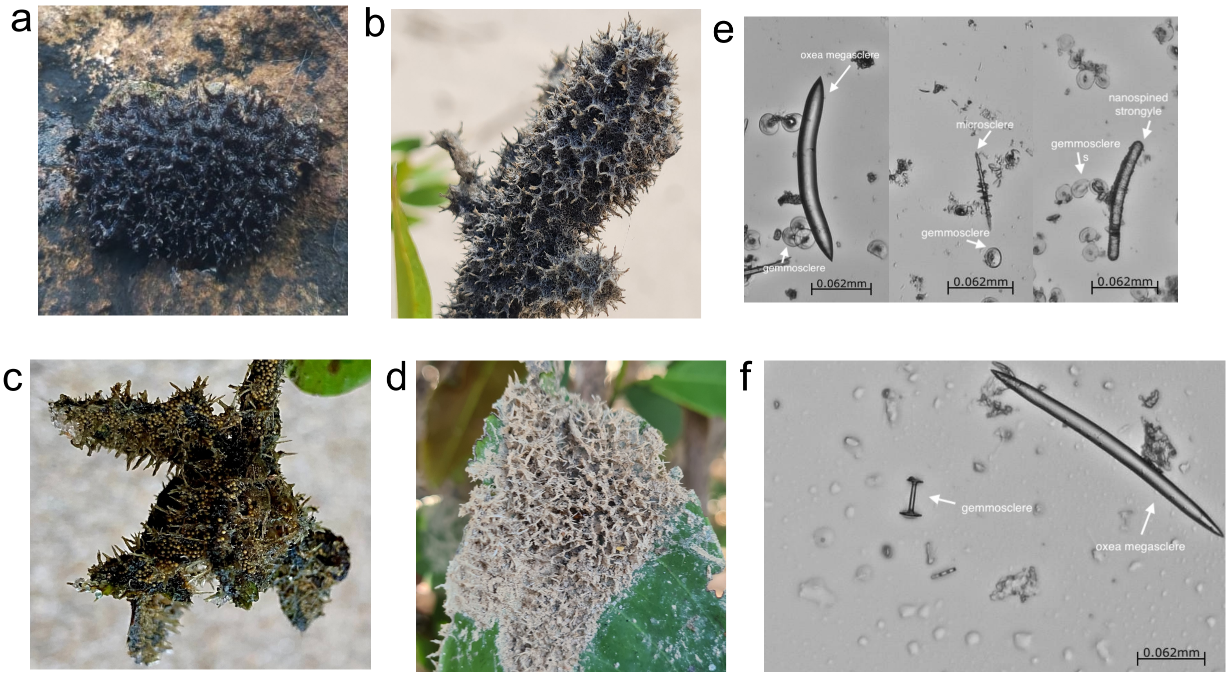


**Figure S1. Amazon freshwater sponges and photomicrograph images of spicules**. The images show both here studied Amazon sponges, *Drulia brownii* (a-b) and *Tubella paulula* (c-d) of non-urbanized sites during the rainy season (a, c), when they are metabolically active and have access to the Tapajós river water, and during the dry season (b, d), without any access to water, with dried and sandy texture; images also show the main morphological traits which allow their identification through morphology, spicules: for *D. brownii* (e), smooth oxeas with abruptly pointed extremities, small, straight and spiny microscleres, and umbonate parmuliform gemmoscleres; for *T. paulula* (f) smooth oxeas, with no microscleres identified and birotuled gemmoscleres. Microscopy magnification is 10x.

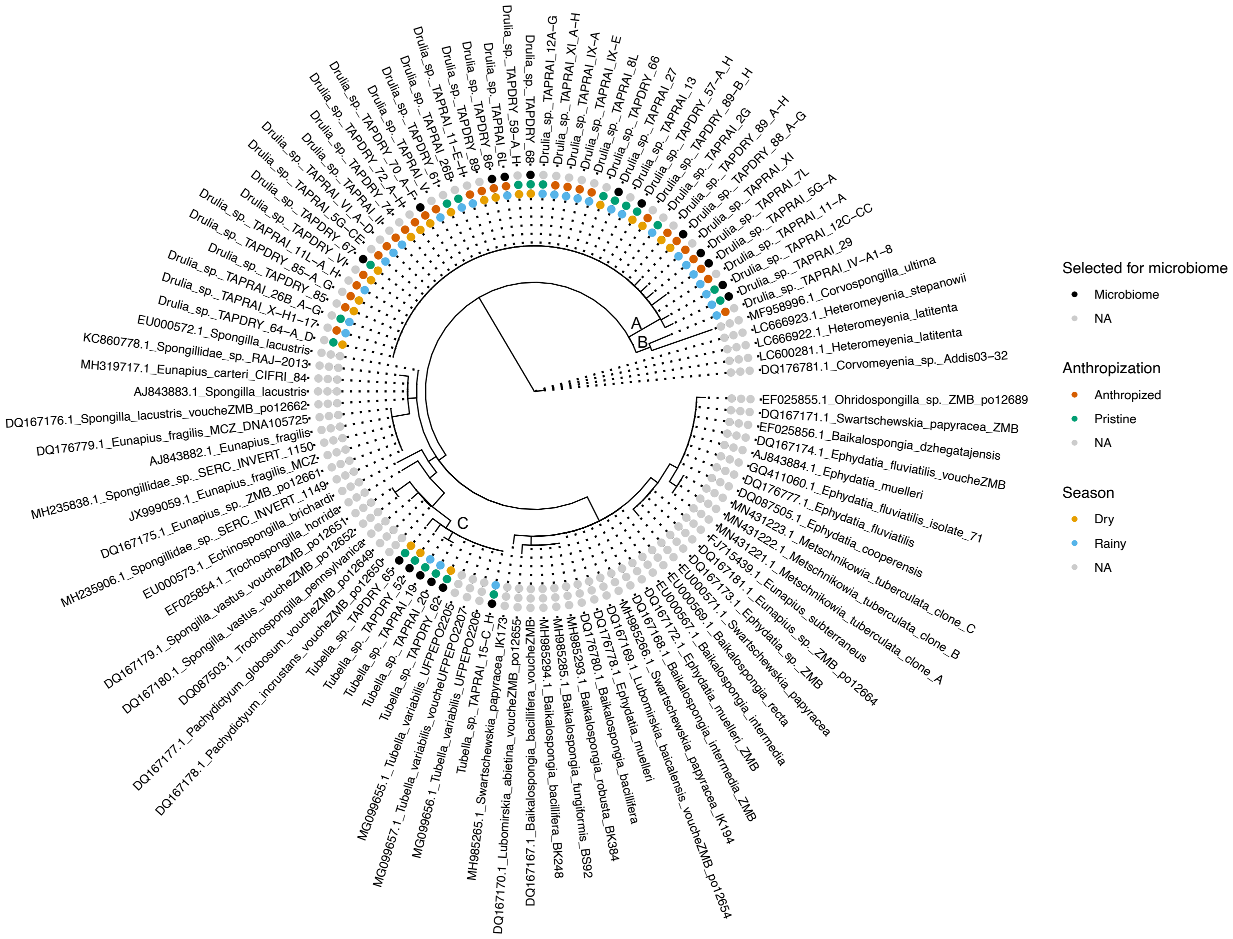


**Figure S2. Amazon freshwater sponge phylogeny.** Circular phylogenetic tree of freshwater sponge samples rooted with *Corvomeyenia* sequences (outgroup). Concentric annotation in layers indicates metadata: outer ring = season (dry or rainy), middle ring = urbanization status (non-urbanized or urbanized), and inner ring = selection for microbiome analysis. The phylogenetic tree was based on the COI partial gene and built using the maximum likelihood method with the GTR+G model with 1000 bootstraps.

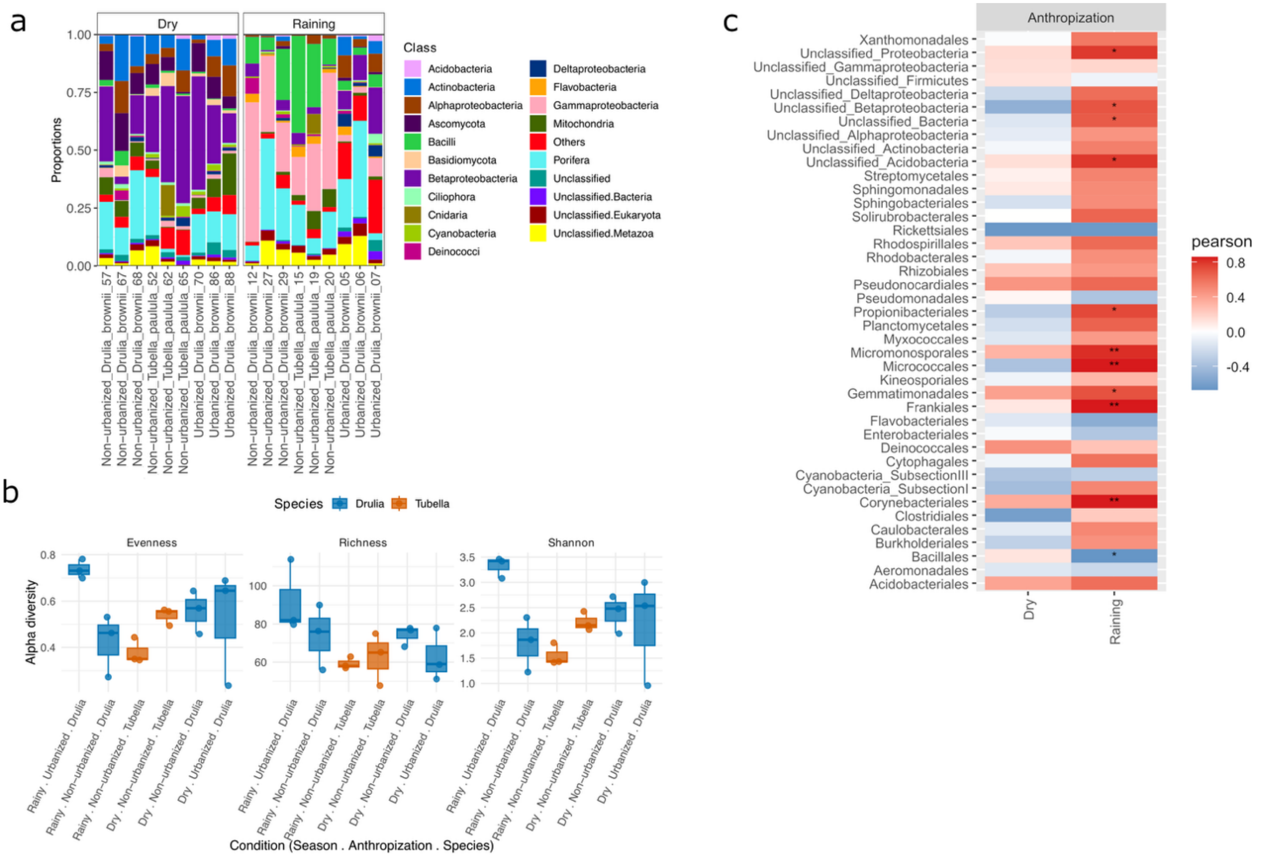


**Figure S3. Whole microbial community of Amazon freshwater sponges across seasons.** The figure (a) shows the relative abundance of the 20 main taxa of the whole metagenome (prokaryotic and eukaryotic) at the class level across sponge samples grouped per condition; (b) box plots with the alpha diversity metrics of sponge-associated microbial orders per sample across conditions. Facets display Richness (observed number of orders), Shannon (H′), and Evenness (Pielou's J′ = H′/ln S). Points are individual samples; boxes show median and IQR; whiskers extend to 1.5×IQR. Colors denote host species: *Drulia brownii* (blue) and *Tubella paulula* (orange); (c) correlations between the 41 most abundant taxa and the variable urbanization under dry and rainy seasons. The colors indicate Pearson correlation direction and strength, with blue indicating negative, red for positive, and asterisks showing statistically significant correlations after FDR correction ($p < 0.05$, $*p < 0.01$, $**p < 0.001$).

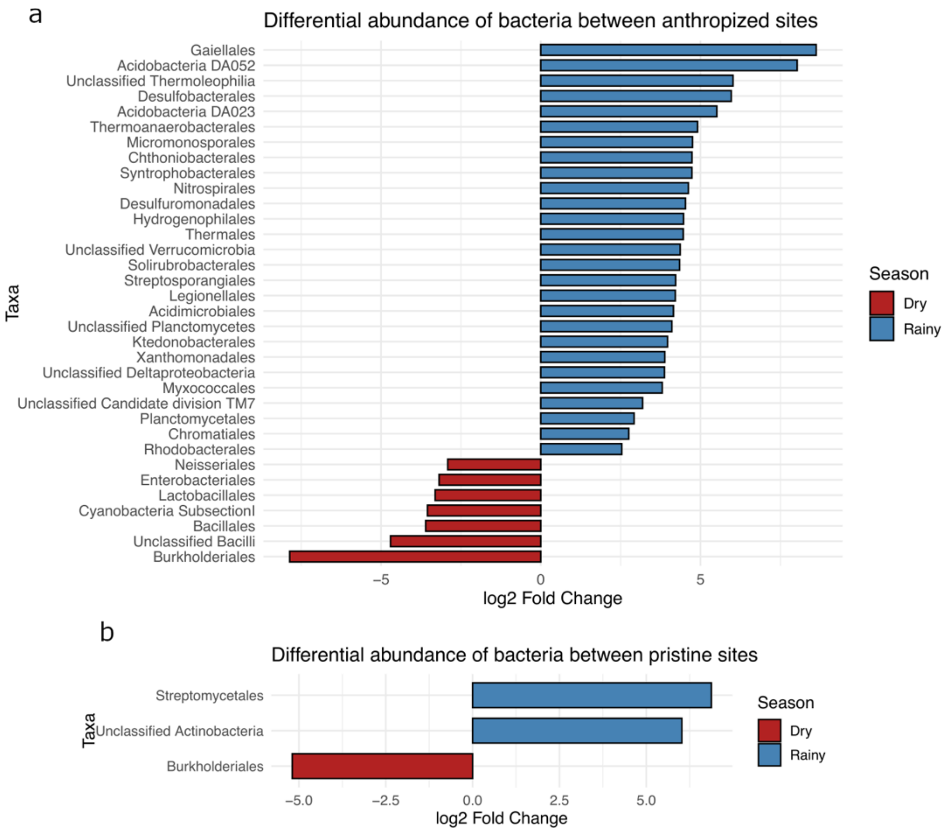


**Figure S4. Gemmule bacterial abundance changes over seasons and sites.** Figures (b-c) show differential abundance analysis of bacterial taxa between seasons in non-urbanized (c) and urbanized (b) sites. Bar plots show taxa with significant differences in relative abundance (DESeq2, padj < 0.05) between the rainy and dry seasons. Positive log2 fold-change values (blue) indicate enrichment during the rainy season, while negative values (red) indicate enrichment during the dry season.

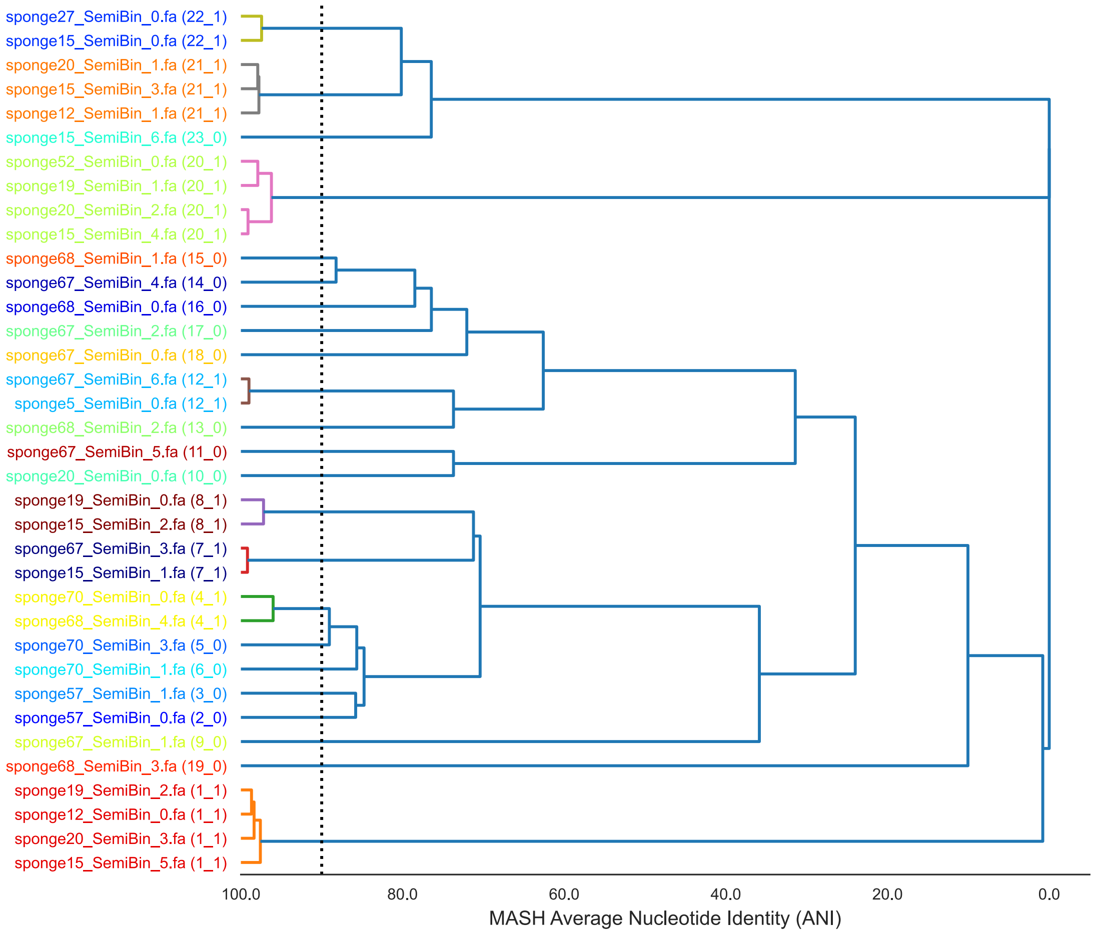


**Figure S5. Metagenome-Assembled Genomes retrieved from whole sponge metagenomes.** The dendrogram represents the clustering of 43 MAGs recovered from freshwater sponge metagenomes. Genomes were dereplicated using dRep through pairwise comparisons based on MASH average nucleotide identity (ANI) to assess genomic relatedness. The x-axis indicates MASH ANI distance, with values closer to 100 representing higher similarity between genomes. The dashed vertical line marks the dereplication threshold of 95% ANI, a commonly used cutoff for delineating species-level clusters. Each tip of the dendrogram corresponds to one MAG, with labels indicating the sample of origin and bin identifier (e.g., sponge27_SemBin_0.fa). Colored branches denote distinct clusters of highly similar genomes, collapsed by dRep according to the ANI threshold. Genomes connected within a branch to the left of the dashed line are considered redundant at the species level and therefore clustered together, while those remaining to the right represent distinct species-level MAGs.

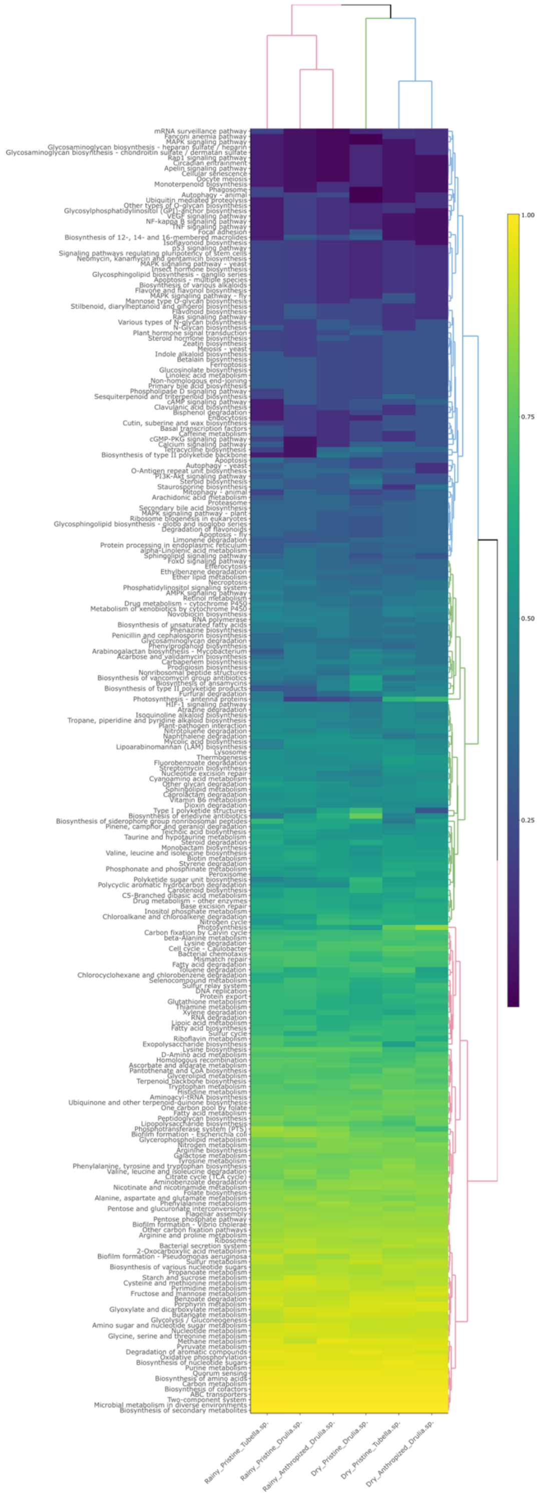


**Figure S6. Whole metagenome assembly KEGG Metabolic Pathways.** The heatmap displays normalized abundance values of KEGG pathways across different assembled metagenomes, based on contigs obtained from the assembly of all three samples representing each site. Colors represent relative abundance levels, with clustering applied to highlight functional similarities between samples and categories. Normalization and percentization were applied to improve comparability.

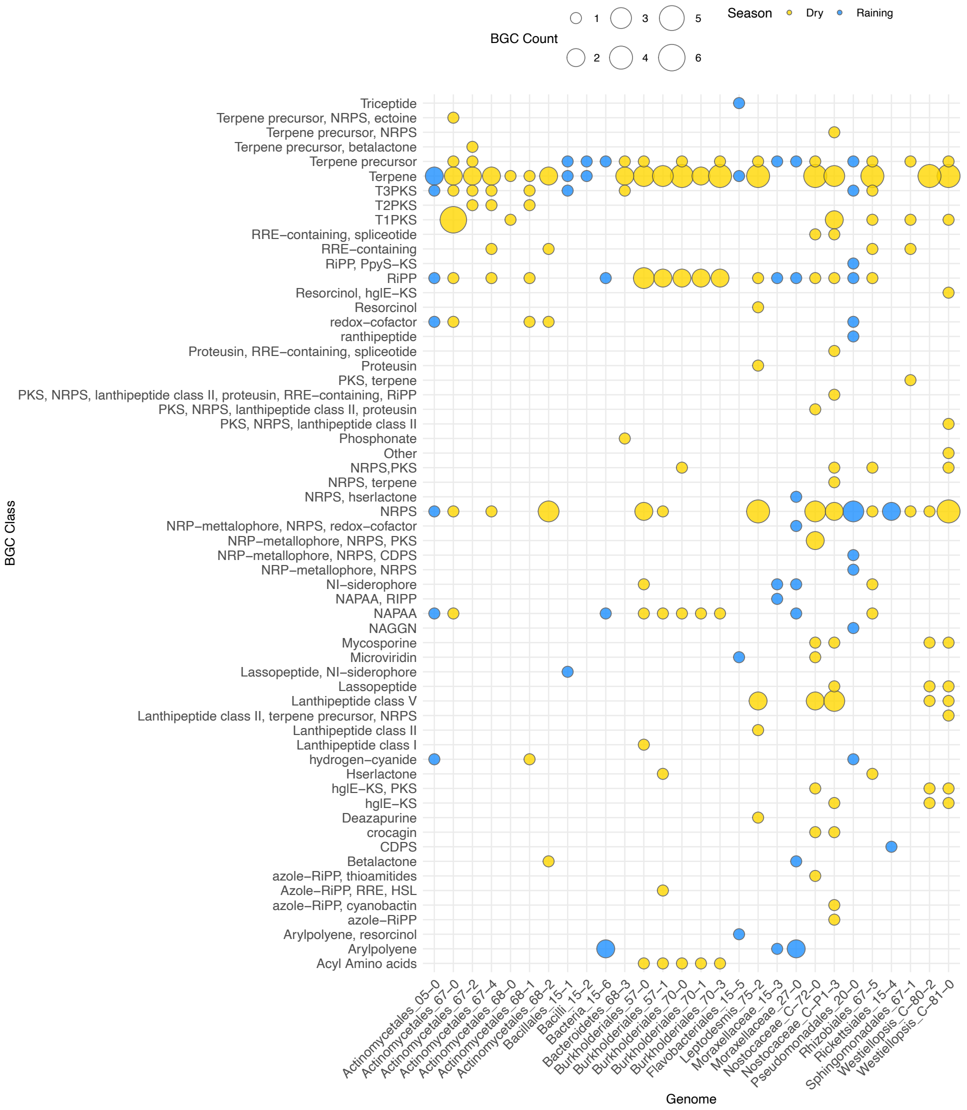


**Figure S7. BGCs identified per genome.** Rows are BGC types and columns are genomes; bubble size scales with the number of BGCs detected in each genome-type cell. Color encodes season (Dry = yellow; Raining = blue). Empty cells indicate no BGCs of that class were found in that genome.

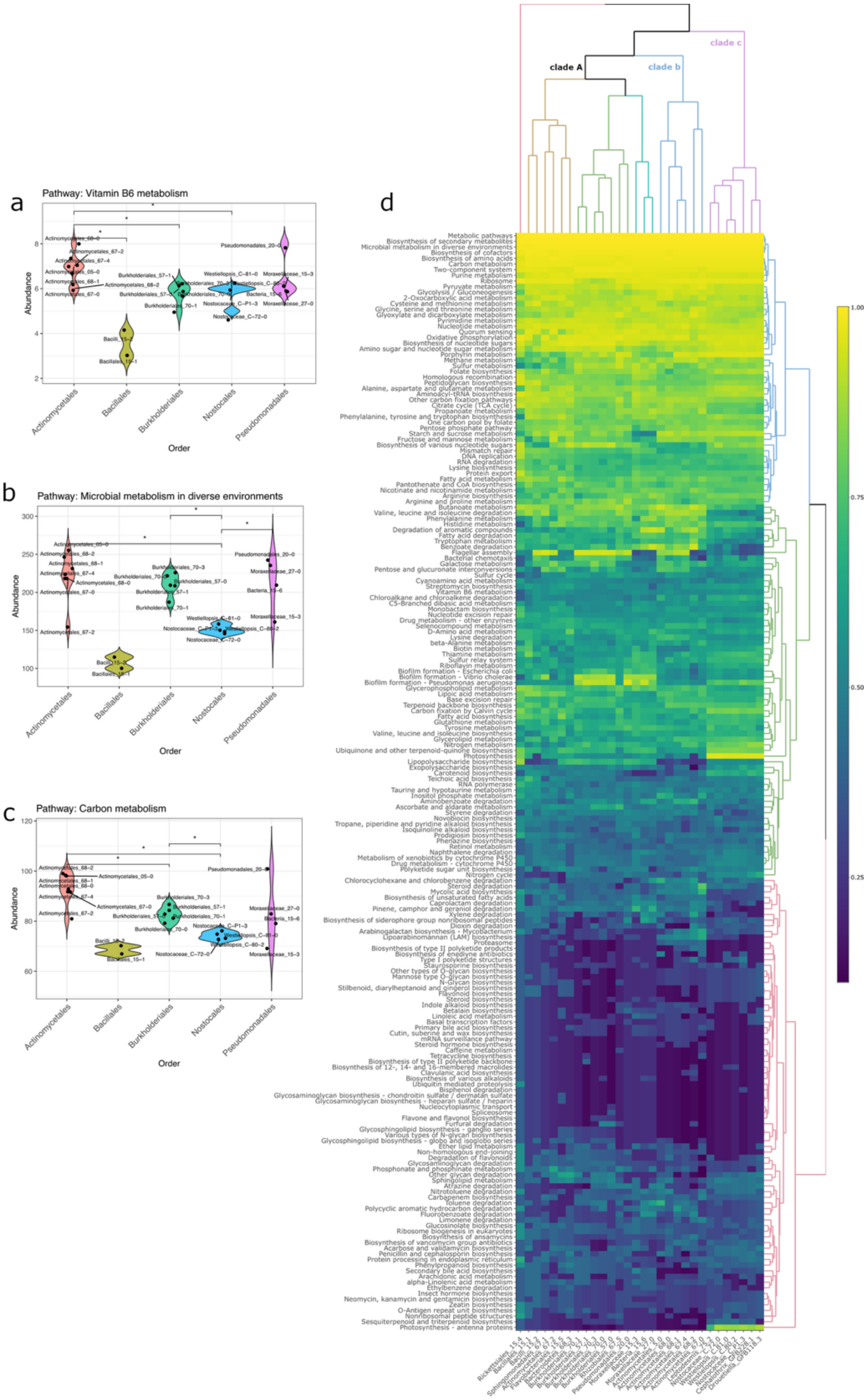


**Figure S8. KEGG Pathways categories across MAGs.** The figure shows (d) a heatmap displaying normalized abundance values of KEGG pathways across different MAGs. Colors represent relative abundance levels, with clustering applied to highlight functional similarities between samples and categories. Normalization and percentization were applied to improve comparability between samples.

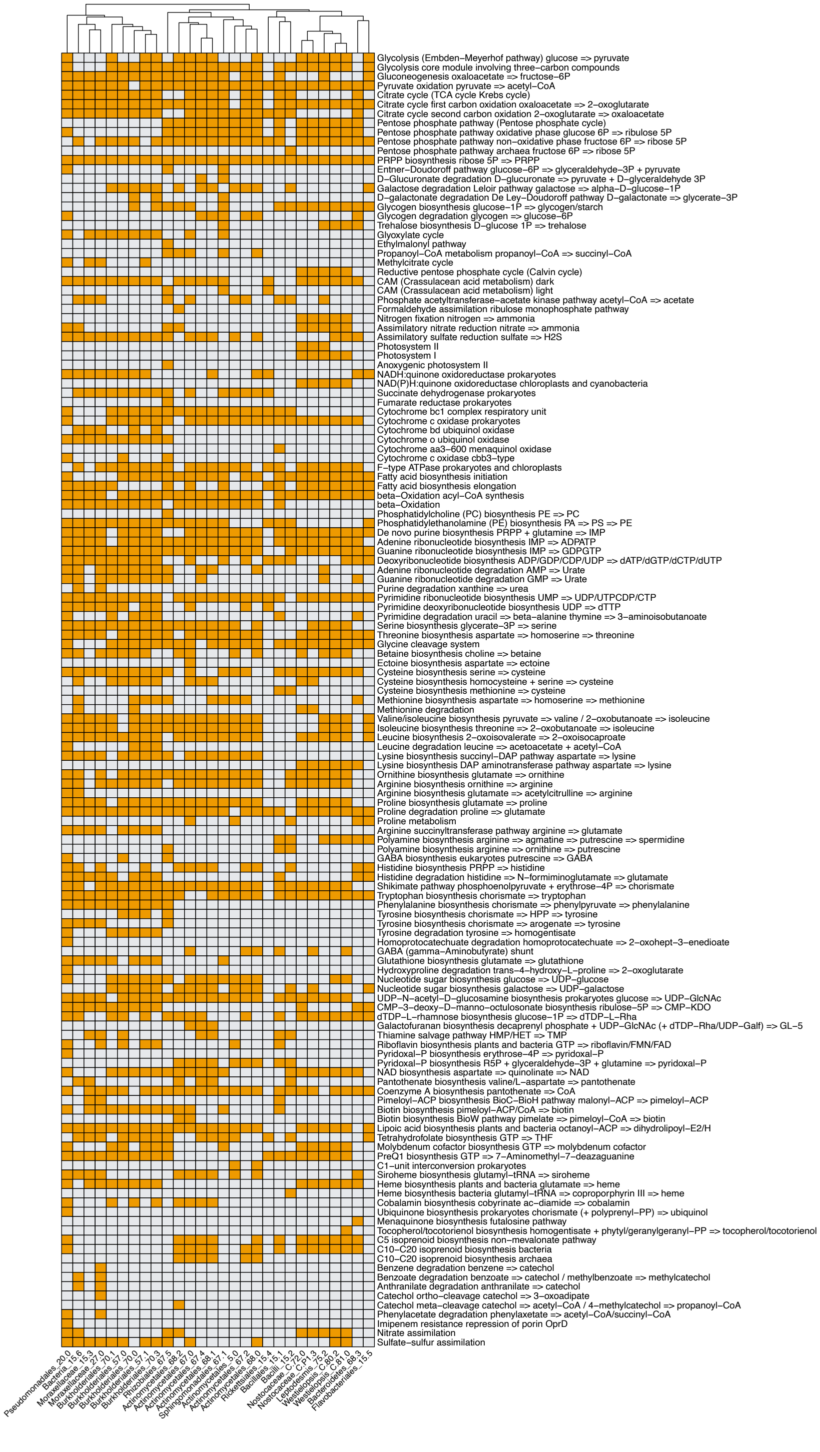


**Figure S9. Complete KEGG modules across MAGs.** Each row represents a KEGG module, and each column corresponds to one MAG. Orange cells indicate the presence of a complete module, while grey cells indicate its absence. Columns (MAGs) are clustered to highlight similarities in metabolic potential.

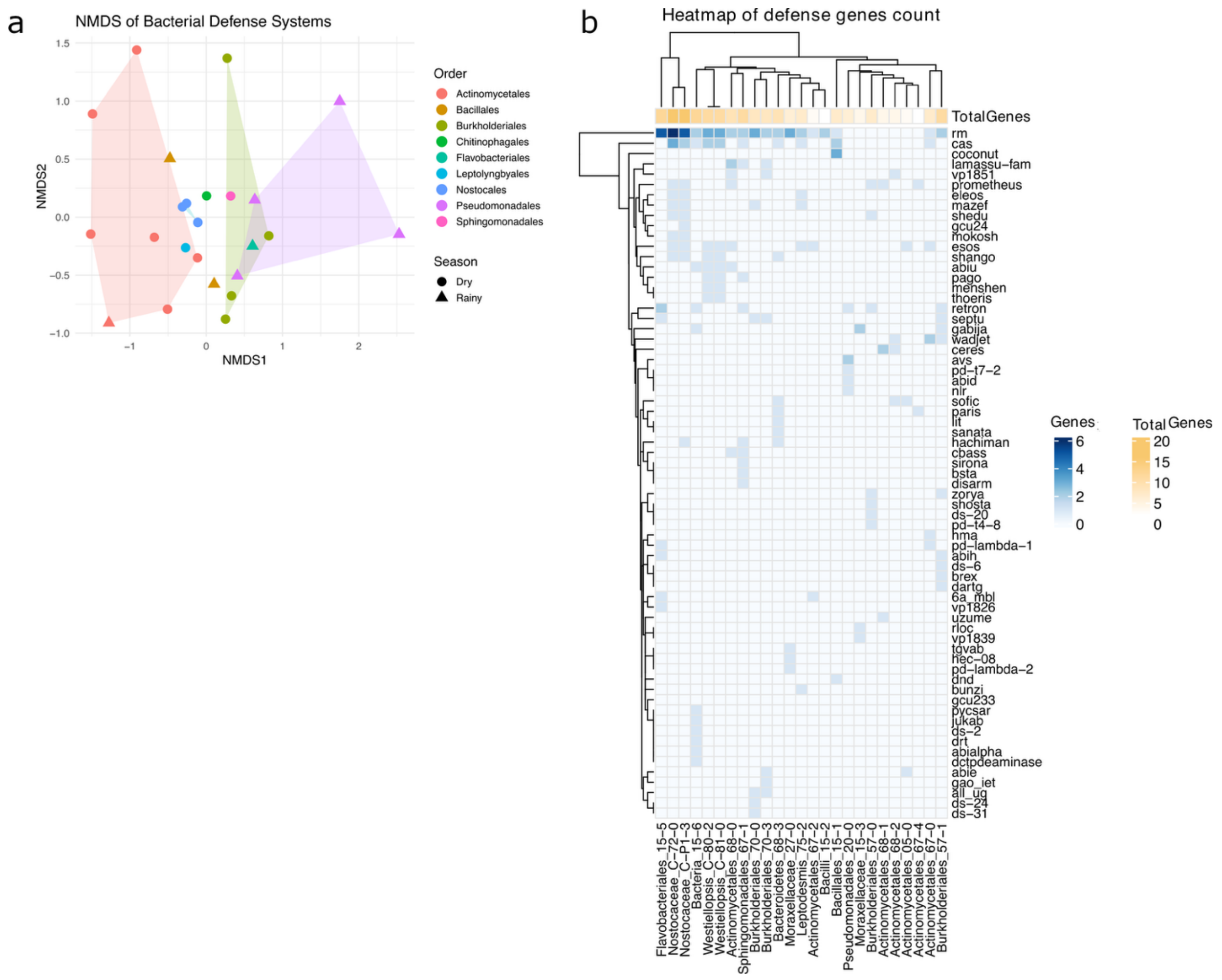


**Figure S10. Defense system genes across freshwater sponge-associated bacterial MAGs.** The figure (a) shows the counts of defense system genes per genome as a heatmap (blue). Rows are the defense categories, and columns are the genomes. Genomes with a total count of zero were removed prior to plotting. Columns and rows were hierarchically clustered to highlight patterns of similarity. A sidebar annotation indicates the sum of defense genes detected in each genome and is displayed at the top (yellow); (b) NMDS based on Bray-Curtis dissimilarities of defense gene profiles. Points represent genomes; colors denote taxonomic Order and shapes indicate Season (Dry = circles, Rainy = triangles). Light polygons are convex hulls drawn only for Orders represented by ≥3 genomes to aid group visualization. Genomes with zero totals were excluded.

**Appendix S1. Isolation and genome sequencing of cyanobacteria from freshwater sponges.**

Freshwater sponge biomass collected during the dry season was used as starting material for microbial isolation. Sponge fragments were inoculated into BG11 medium (with nitrogen) and $BG11_0$ medium (nitrogen-free) and incubated at room temperature (approximately 30 ± 3 °C) under a photoperiod of 16 h light and 8 h dark, simulating natural local light conditions. While BG11 cultures were enriched mainly with eukaryotic algae and showed limited cyanobacterial proliferation, $BG11_0$ cultures favored cyanobacterial growth and provided the most successful enrichment for isolation. Cyanobacteria were subsequently isolated by serial “fishing” on Kline Concavity Slides using dilutions of $10^{-1}$, $10^{-2}$, $10^{-3}$, and $10^{-4}$, and unialgal cyanobacterial isolates were recovered. Biomass from pure cultures was harvested, and genomic DNA was extracted using the FastDNA SPIN Kit for Soil (MP Biomedicals) according to the manufacturer’s instructions. DNA was quantified and assessed for purity prior to sequencing. Nanopore sequencing was performed on the PromethION P2 Solo platform using the Rapid Sequencing Kit. The workflow included end-prep and transposase fragmentation of high-molecular-weight DNA, adapter ligation, and loading onto R10.4.1 flow cells. Sequencing output was basecalled using Dorado with high-accuracy (HAC) models, followed by quality filtering ($Q \geq 12$). De novo assembly was performed with Flye v2.9 in metagenome mode. Genome binning was conducted using SemiBin2 with default parameters. The resulting metagenome-assembled genomes (MAGs) were assessed for completeness and contamination with CheckM2.

### Appendix S2. Sampling and cultivation strategy to obtain *Pseudomonas* strains

Sampling was carried out during the rainy season, in which freshwater sponges were collected from a non-urbanized region in order to obtain samples for cultivation and isolation of *Pseudomonas* strains. After collection, the samples were refrigerated at -8 °C and transported to the Laboratory of Bacteriology of the Universidade Federal do Oeste do Pará (LaBac - UFOPA) for processing. Samples were macerated using a sterile mortar and pestle and homogenized in a 0.85% NaCl solution. A pre-enrichment step was carried out by mixing the homogenate at a 1:1 ratio (sample:enrichment medium) with Brain Heart Infusion (BHI, Kasvi) broth supplemented with 10% bidistilled glycerol, to promote the growth of *Pseudomonas* spp. The cultures were incubated in a shaker for 24 h. Serial dilutions were then prepared at a 1:9 ratio using 0.85% NaCl (1 mL inoculum into 9 mL NaCl) up to the fifth dilution ($10^{-1}$ to $10^{-5}$). From each dilution, 100 µL were plated onto Pseudomonas Isolation Agar (PIA, Himedia) using the spread plate technique with sterile Drigalski loops. Plates were incubated for 24 h at 35 ± 1 °C in a bacteriological incubator. After incubation, colonies that grew on PIA were selected, isolated, and transferred with a sterile loop onto Tryptone Soy Agar (TSA) slants, followed by incubation under the same conditions described above. For characterization and confirmation of the target bacterium, isolates were subjected to biochemical tests. Gram staining followed by light microscopy was first performed to verify whether the isolates were Gram-negative rods. Subsequent assays included oxidase activity, nitrate reduction, motility, glucose fermentation, and growth on Cetrimide Agar, to confirm the presence of *Pseudomonas aeruginosa*. Pure cultures were maintained on TSA for subsequent analyses. For molecular identification, colonies were suspended in 50 µL of ultrapure water, boiled at 50 °C for 10 minutes, and then centrifuged at 13,000 rpm for 10 minutes. Five microliters of the supernatant were used as template for PCR amplification of the 16S rRNA gene with the universal reverse primer. Sequencing was performed using Sanger sequencing with the 3500 Genetic Analyzer (Applied Biosystems) and BigDye Terminator v3.1 chemistry (Thermo Fisher). Sequence data were analyzed using Sequencher and BioEdit software, and taxonomic identification of the isolates was performed through BLAST searches against the NCBI database.

### SUPPLEMENTARY TABLE LEGENDS

**Table S1.** List of freshwater sponge sequences retrieved from GenBank and used for phylogenetic reconstruction. The table provides the GenBank accession number and corresponding taxonomic identification (species or voucher/clone designation) of each sequence included in the analysis. A total of 55 sequences representing mainly lineages of Spongillidae were used to infer the phylogeny.

**Table S2.** Quality assessment and taxonomic classification of MAGs recovered from the metagenomes. A total of 43 MAGs with ≥70% completeness and ≤10% contamination were retained after initial filtering. Dereplication was performed using dRep, and genome quality and taxonomy were evaluated with CheckM. The table shows the CheckM marker lineage, estimated completeness, and contamination for each MAG; the ones marked with an asterisk (*) represent the best dereplicated representatives and were selected for downstream analyses.

**Table S3.** Summary of predicted biosynthetic gene clusters (BGCs) identified in MAGs and respective detailed information. Summary and corresponding sheets for bacterial MAGs containing the following information for all detected BGCs: BGC name, BGC type, BGC length (bp), most similar known cluster, and confidence score as predicted by antiSMASH.
